# Chronostratigraphy of Jerzmanowician. New data from Koziarnia Cave, Poland

**DOI:** 10.1101/2020.04.29.067967

**Authors:** Małgorzata Kot, Maciej T. Krajcarz, Magdalena Moskal-del Hoyo, Natalia Gryczewska, Michał Wojenka, Katarzyna Pyżewicz, Virginie Sinet-Mathiot, Marcin Diakowski, Stanisław Fedorowicz, Michał Gąsiorowski, Adrian Marciszak, Paweł Mackiewicz

## Abstract

Lincombian-Ranisian-Jerzmanowician (LRJ) sites are sparse, and Koziarnia Cave in Poland is one of only few such sites situated at the eastern fringe of LRJ. The aim of the recent study was to obtain new chronostratigraphic data for the LRJ industries due to their extreme scarcity in Central Europe. Although the new fieldworks did not bring new *fossil directeur* such as bifacial leafpoints, a detail debitage analysis enabled identifying a presence of the ventral thinning chips in layer D, which could be identified as the LRJ assemblage-containing stratum. Besides the LRJ assemblage, strata with traces of Late Middle Palaeolithic and Early Gravettian occupation were found at the site. The radiocarbon dates of Koziarnia samples show that the archaeological settlement represent one of the oldest Gravettian stays north to Carpathians. What is more, these dates demonstrate that the cave had been alternately occupied by humans and cave bears. Additionally the radiocarbon dates indicate rather young chronology of the Jerzmanowician occupation in Koziarnia Cave (c.a. 39-36 ky cal. BP). The results confirm the possibility of long chronology of the LRJ technocomplex, exceeding the Campanian Ignimbrite event.

## Introduction

Middle/Upper Palaeolithic transitional industries in Central Europe are among the most ephemeral and most debated topics in Palaeolithic discourse [1–3]. After over 100 years of research into the subject, we are still seeking for answers to crucial questions regarding, e.g. the origins [4–8], the chronology [9, 10], internal divisions [11–13], or even the identification of the population responsible for these industries [14–19].

Lincombian-Ranisian-Jerzmanowician is one of such transitional industries determined by the presence of bifacially worked leafpoints made on blades obtained from double platform cores. Technological and experimental studies show that one is dealing here with a predetermined technique of obtaining leaf-shaped blades, which were later adjusted to a minimal extent to mirror the exact willow leaf shape through ventral thinning. Such technological features are present in transitional assemblages in southern Poland (Nietoperzowa Cave), Southern Germany (Ranis), Belgium (Spy) and the southern part of Great Britain (Beedings, Kent’s Cavern), but are somewhat absent to the south of the Carpathians, where a Szeletian type of industry prevails [14, 20–23].

The term “Jerzmanowician” was used for the first time in 1961 by W. Chmielewski after his studies in Nietoperzowa Cave located in Jerzmanowice village [20]. Chmielewski focused his research on two cave sites, Koziarnia and Nietoperzowa, where the first bifacial leafpoints were found already in the 19^th^ century.

In the second half of the 19^th^ century, the cave sediments of several sites were exploited by local landlords to be sold as field fertiliser. The southern part of the Polish Jura, a karstic region rich in caves, was at that time the part of the Russian Empire, but the business was driven by Prussian businessmen, who organized the transit of train wagons filled with cave sediments to Prussia. In consequence, the sediments of such caves as Nietoperzowa, Koziarnia and Gorenicka were heavily destroyed. Due to the sediment exploitation, the original sediment level in Nietoperzowa Cave was lowered by around 1.5-2 m, whereas in Koziarnia Cave by around 0.5-1 m. During the cave sediment exploitation, multiple prehistoric animal bones and artefacts were found. The discoveries led Ferdinand Römer, a geologist and palaeontologist from Schlesische Friedrich-Wilhelms-Universität in Breslau (now University of Wrocław), to study the cave sediments in detail. For this reason, he collected the already discovered artefacts and conducted his own excavations at several caves in the region. The various findings discovered by F. Römer [24, 25] include bifacial leafpoints from Nietoperzowa and Koziarnia caves.

To check their stratigraphy, Chmielewski re-excavated both caves in the 1950s. In Nietoperzowa Cave, he found three archaeological horizons (layers 4, 5a and 6), containing in total more than 87 bifacially worked leafpoints and their fragments [26]. In Koziarnia Cave, he opened ten trenches covering the area of 120 m^2^ in total, but none of the 21 layers determined inside the cave and on the terrace in front of the cave could be clearly described as containing the Jerzmanowician assemblage. One of the layers, i.e. 13, which he claimed did not contain any stone artefacts, was black in colour due to the high amount of charcoal [27]. Chmielewski called it a “cultural layer”, and by comparing it to layers 4 and 6 in Nietoperzowa Cave, he initially suggested that it was a Jerzmanowician horizon [20]. No radiometric dates were obtained at that time to confirm the hypothesis.

Even though the determination of the Jerzmanowician culture was based mostly on the Nietoperzowa Cave assemblage, the Koziarnia and Mamutowa caves were also included. A single radiocarbon date (38 160 ± 1250 BP, Gro-2181) obtained for a wood charcoal from layer 6 in Nietoperzowa Cave was presented by Chmielewski in 1961 [20], and since then it has been treated as the major chronological framework of the whole technocomplex. It was later proposed that all the assemblages containing bifacially-worked blade leafpoints from the European Plains can be merged into one category – Lincombian-Ranisian-Jerzmanowician (LRJ), a term widely used till today [28].

The chronology of Jerzmanowician has been restudied several times since then [14, 29, 30, 32]. The analyses were in most cases conducted on the animal fossil collection. The most recent results of multiple radiocarbon dating obtained on the basis of cave bear remains from Nietoperzowa Cave [33] showed the limitation of the possible use of the old collection. The radiocarbon range of each stratum shows all the chronological spectra observed in the cave, which might indicate problems resulting from the exploration, documentation or mixing of the collection. Only a new detailed fieldwork would help to resolve all the chronological issues linked to LRJ industries.

In order to clarify the chronostratigraphic position of Jerzmanowician, a new fieldwork project was initiated in 2017. It aimed at the verification of the stratigraphy of Koziarnia Cave and obtaining reliable radiometric dates for a complete profile of the site, as well as reconstructing the palaeoenvironmental conditions for particular strata [34]. The paper presents the obtained chronostratigraphic data with a comparison to the results published before.

### Koziarnia Cave

Koziarnia Cave is located in Sąspów Valley, in the southern part of the Polish Jura (Fig 1). The cave has a 5-metre-high entrance heading SW with the main chamber covering an area of over 100 m^2^ behind it and a single 40-metre-long gallery narrowing toward the end of the cave.

**Fig 1.**
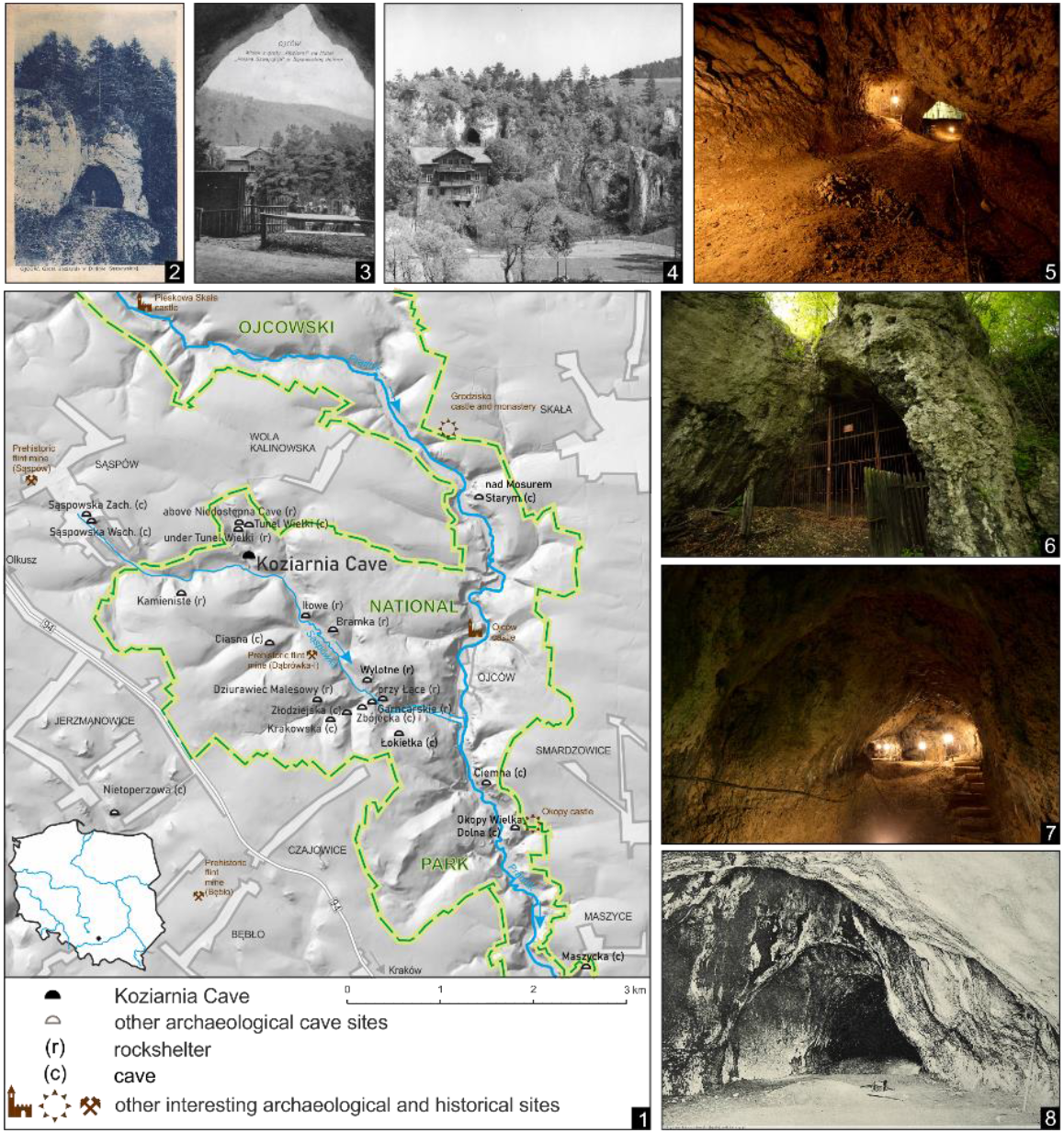
Koziarnia Cave, its surroundings and state of preservation. (1) Localisation of Koziarnia Cave. (2) Postcard dated to 1927 illustrating the entrance to Koziarnia Cave. (3) 1^st^ half of the XXth century, view from the Koziarnia cave on the “Szwajcaria” hotel, situated on the opposite slope of Koziarnia Gorge. At that time a dance floor was built inside the cave. (4) Koziarnia Cave and Villa Koziarnia (previously “Szwajcaria” hotel) during excavations of prof. Chmielewski in 1958-1962 (photo from the archives of prof. T. Madeyska-Niklewska). (5-6) Koziarnia Cave during excavations in 2017. (7) Current state of preservation of Koziarnia cave sediment. The sediments inside of the cave are partly destroyed by collapsed unfilled archaeological trenches and ditches made during installation of the seismograph at the back of the cave. (8) State of preservation of Koziarnia Cave sediment in 1910. The pit visible in the middle part may be a remnant of the F. Römer fieldworks in 1879.

The cave was continuously in use until World War I. At the beginning of the 20^th^ century, when the sediment exploitation was halted, a dance floor was built in the main chamber, and a resting place was located on the terrace in the front of the entrance. In 1919, the cave was excavated by S. Krukowski, who at the same time conducted fieldwork in the nearby Ciemna Cave [35, 36]. Krukowski made a trench in the entrance zone of the cave, finding nothing but some Holocene artefacts, which he never published. In 1958-62, the cave was excavated again by W. Chmielewski. He found most of the sediments in the main chamber already destroyed due to the previous activities. Inside the cave, undisturbed layers were found as far as 20 m from the cave entrance. In his final publication, Chmielewski described the cross-section as having 21 separate geological layers [27]. Several of them contained flint artefacts. Originally, Chmielewski distinguished eight cultural layers (4, 7, 10, 13, 16B, 17, 18, 20), out of which layers 4, 7, 10 and 13 contained only charcoal and no lithic artefacts.

Lithic artefacts were found only in the lower layers. Middle Palaeolithic settlement was associated with layers 17, 18 and 20. The small assemblage contains two bifacial backed knives, several flake discoid cores and post-depositionally damaged debitage. Rare stone artefacts also described as Middle Palaeolithic were found in layers 15 and 16. One of the artefacts found in layer 15 (mistakenly published as coming from layer 17) is a massive blade made on a double platform core with multiple post-depositional retouches (Fig 7, IX/17-23/61). The archaeologically sterile layer 13 was described as black due to high charcoal concentration. The concentration of charcoal was very high in the main corridor (20–35 metres from the entrance). The upper part of the section was present only to a minimal extent in the trenches located farther from the entrance. Chmielewski correlated them with the loess sequences found in the front of the cave and dated to MIS 2.

**Fig 7.**
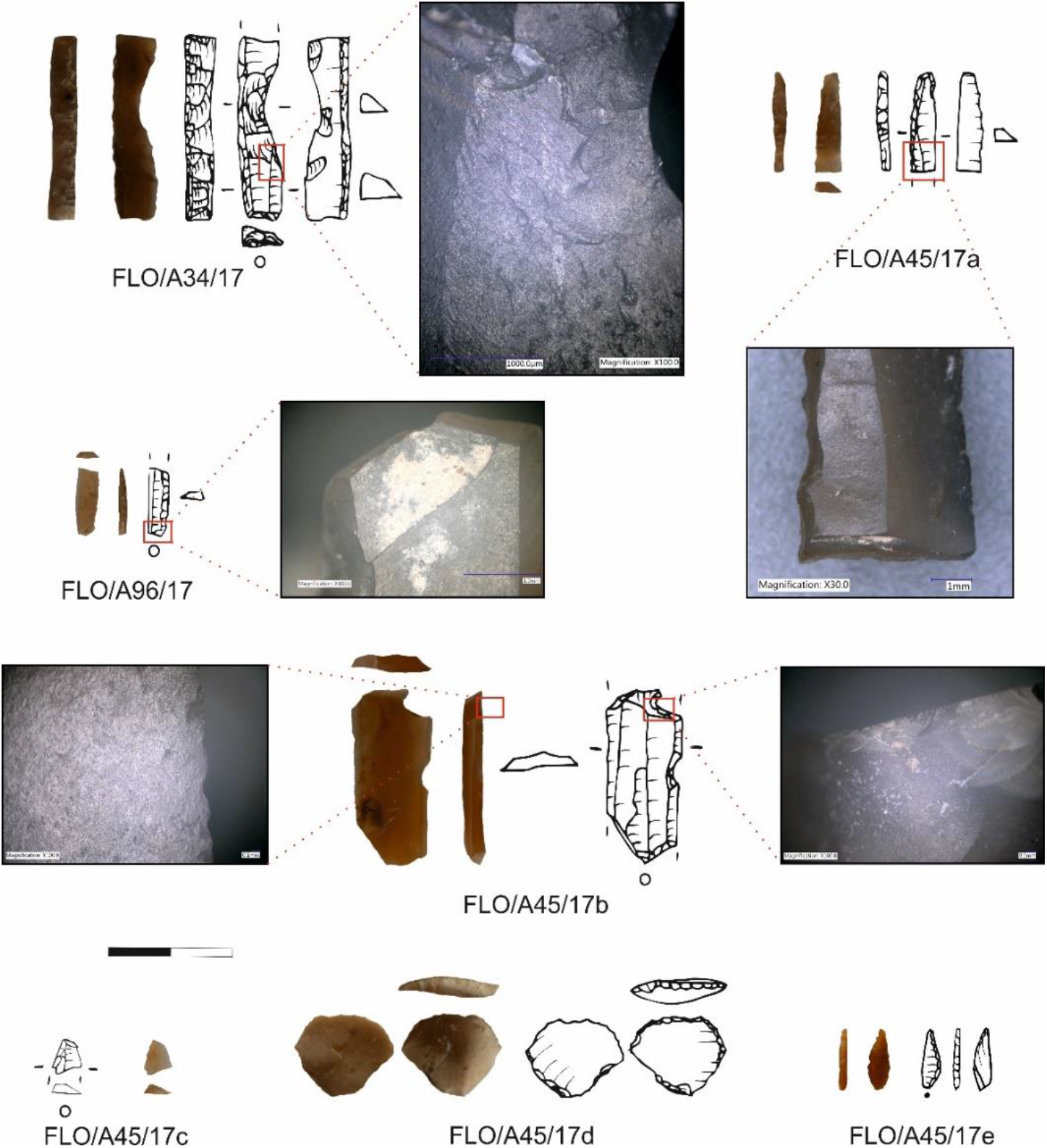
Gravettian artefacts found in layer K and K’ in trench IX/2017.

Unfortunately, the trenches were not refilled after the excavations, and they have stayed open for 70 years. All the walls disintegrated slowly causing massive damage (Fig 1.7).

## Methods & Materials

The new fieldwork conducted in 2017 covered 2.85 m^2^ (Fig 2). A trench was opened 40 metres from the cave entrance in the NW corner of Chmielewski’s trench IX in order to correlate the stratigraphy and open a section in the place that could cover the highest possibly undisturbed profile. The collapsed walls of the old trenches containing a mixed sediment were visible during the fieldwork. One should still take into consideration that even the stratified parts of the trench might be post-depositionally moved due to the slight trench wall movements. The *in situ* sediment was collected and wet sieved in whole with 1 mm sieves. A mixed sediment was collected and wet sieved with 3 mm sieves. The sieved material was dried and screened in order to collect tiny microfaunal, anthracological and archaeological material. All the *in situ* findings, including the charcoal, were 3D measured.

**Fig 2.**
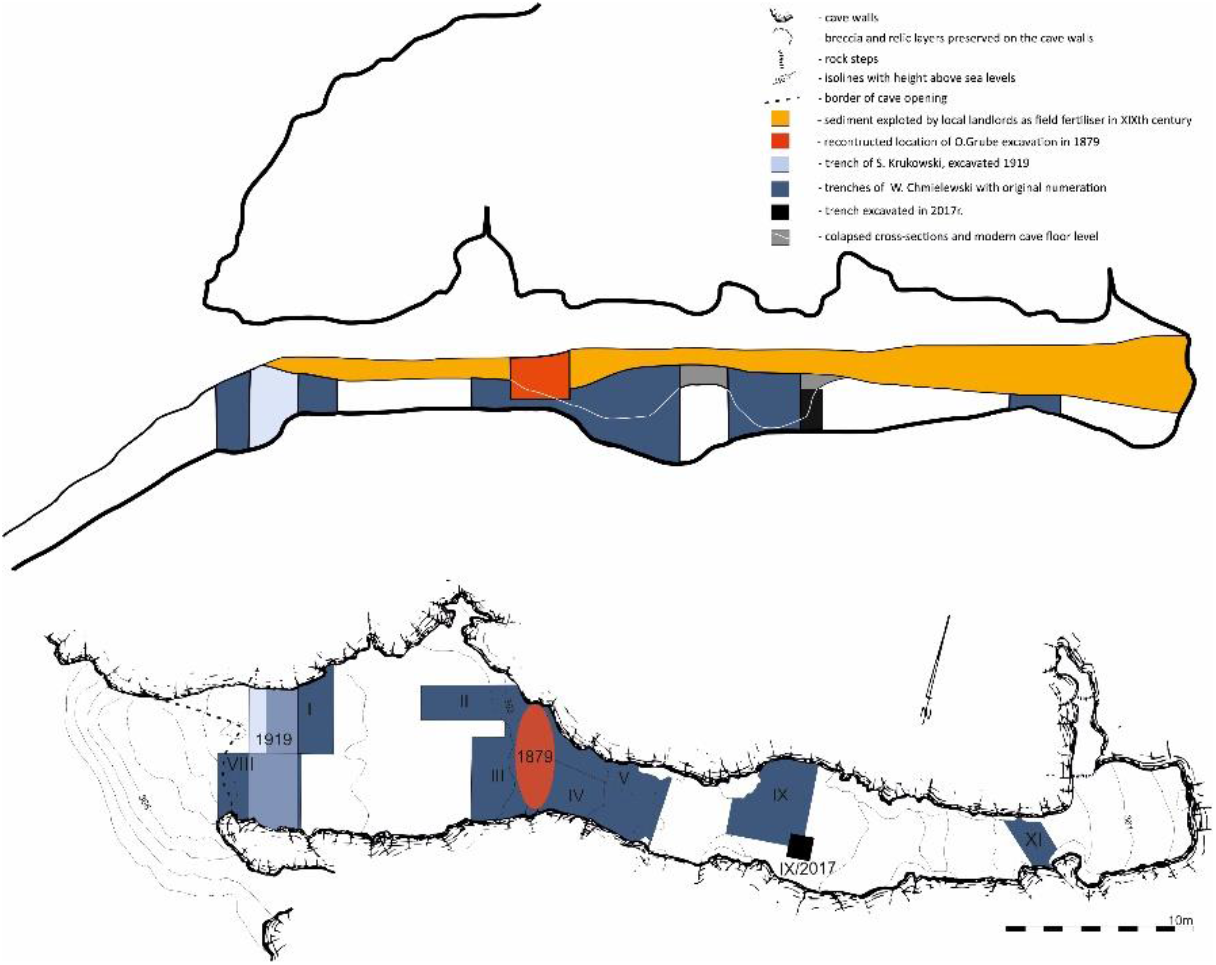
Plan and the cross-section of Koziarnia Cave with the localisation of all previous archaeological fieldworks.

Additionally, the old collection of artefacts found in Koziarnia by F. Römer in 1879 was restudied. For the purpose of chronostratigraphic studies, two unpublished bone tools were radiocarbon dated and analysed through zooarchaeology by mass spectrometry (ZooMS) and traseology.

### Radiocarbon dating

In each layer, charcoal was the dominated material (Table 1). Each charcoal fragment was taxonomically identified on the basis of wood anatomy atlases [37, 38] and the modern wood collections of the Department of Palaeobotany of the W. Szafer Institute of Botany of the Polish Academy of Sciences. Only identified fragments were dated. The selection of the most suitable charcoal fragments is based on a list of woody flora typical of the environmental conditions for a specific period, as well as the size and ring curvature indicating the origins of the wood from a branch or a trunk in order to avoid the old wood problem [39, 40]. For MIS 3, it is more adequate to select taxa representing coniferous wood better adapted to the colder conditions of the Pleistocene in Central Europe [41–45]. When no charcoal fragments were available from a chosen stratum, animal fossils were used for dating. Due to the proximity to the edge of the old trenches and possible contamination of the sediment, only samples collected in the farthest part from the edge of the old trench were used.

**Table 1.** Radiocarbon dates from Koziarnia cave.

| date code | sample inv. No | dated material | species | trench | original layer | final layer | C14 date (uncalibrated) | C:N ratio in bone collagen | comments |
| --- | --- | --- | --- | --- | --- | --- | --- | --- | --- |
| OxA-39509 | MMW/26 61.4 | bone tool | *Elephantidae* | ?/1879 | ? | ? | 22020 ± 150 BP | - |  |
| OxA-39539 | MMW/26 61.5 | bone tool | *Elephantidae* | ?/1879 | ? | ? | 20870 ± 210 BP | - |  |
| Poz-82394 | - | bone | *Ursus arctos priscus* | ?/1879 | ? | ? | 39200 ± 1100BP | 2.94 | 8.4% coll |
| Poz-119319 | KOZ_IX_4_5 | bone/man dible | *Ursus ingressus* | IX/1961 | 4/5 | 3-5 | 25440 ± 210 BP | 3.08 |  |
| Poz-119320 | KOZ_IV_2 | bone/skul 1 | *Ursus ingressus* | IV/1960 | 2a | 9-11 | 27340 ± 260 BP | 3.08 |  |
| GdA-3898 | Koz-04-12 | tooth/M2 | *Ursus ingressus* | IX/1961 | 12? | 10? | 32440 ± 240 BP | 3.5 |  |
| Poz-98895 | C9 | charcoal | *Picea abies/Larix decidua* | IX/2017 | K | 12 | 28090 ± 360 BP | - | 0.3 mgC |
| Poz-98898 | C52 | charcoal | *Pinus sp.* | IX/2017 | K' | 13 | 30330 ± 500 BP | - | 0.2 mgC |
| Poz-99773 | B68 | tooth/I | *Ursus ingressus* | IX/2017 | D | 15 | 37650 ± 900 BP | 3.19 | 1.9% coll |
| Poz-99816 | B254 | tooth/M1 | *Ursus ingressus* | IX/2017 | D | 15 | 39000 ± 1000 BP | 3.17 | 4.0% coll |
| Poz-110657 | C102 | charcoal | *Pinus type sylvestris-mugo* | IX/2017 | F | 16b | 33100 ± 1200 BP | - | 0.14 mgC |
| GdA-3896 | Koz-01-7 | tooth/I3 | *Ursus ingressus* | IV/1960 | 7 | 15-16 | 39340 ± 430 BP | 2.8 |  |
| Poz-98901 | C101 | charcoal | *Pinus type sylvestris-mugo* | IX/2017 | H'-cleaning section | 17 | 33230 ± 480 BP | - | 0.8 mgC |
| Poz-99815 | B165 | tooth | *Ursus ingressus* | IX/2017 | H' | 17 | 40100 ± 1100 BP | 3.17 | 4.3% coll |
| Poz-99814 | B160 | bone/met apodium | *Ursus ingressus* | IX/2017 | H' | 17 | >45000 BP | 3.19 | 4.5% coll |
| Poz-116687 | B200 | tooth | *Ursus ingressus* | IX/2017 | I | 17 | 40600 ± 1200 BP | 3.13 |  |
| GdA-3897 | Koz-02-10_10a | bone/metapodium | *Ursus ingressus* | V/1960 | 10-10a | 19-20 | 24190 ± 120 BP | 3.1 |  |
| Poz-99806 | B125 | bone/phalange | *Ursus ingressus* | IX/2017 | F | 16b | 26160 ± 180 BP | too low collagen quantity | 0.3% coll, 0.9 mgC |
| Poz-98896 | C26 | charcoal | *Picea abies/Larix decidua* | IX/2017 | F | 16b | 29430 ± 720 BP | - | 0.13 mgC; incorrectly carbonised |
| Poz-98425 | C63 | charcoal | *Pinus type sylvestris-mugo* | IX/2017 | D | 15 | 220 ± 30 BP | - |  |
| Poz-98902 | C99 | charcoal | *Juniperus communis* | IX/2017 | E | 16a | 145 ± 30 BP | - |  |
| Poz-98899 | C56 | charcoal | *Pinus type sylvestris-mugo* | IX/2017 | L/M | 19-21 | 140.23 ± 0.37 pMC |  | 0.7 mgC |
| Poz-98900 | C57 | charcoal | *Pinus type sylvestris-mugo* | IX/2017 | M | 21 | 175 ± 30 BP |  | 0.6 mgC |

Charcoal, bones, teeth and ivory were dated with the AMS radiocarbon method in the Poznań Radiocarbon Laboratory (Poland), the Oxford Radiocarbon Unit (UK) and the Gliwice Absolute Dating Methods Centre (Poland). In the case of bone, teeth and ivory artefacts, the dated fraction was collagen, and in the case of charcoal, it was cellulose. Collagen and cellulose extraction and purification followed widely accepted methodology [46, 47].

The obtained radiocarbon ages were calibrated versus the INTCAL’13 radiocarbon atmospheric calibration curve [48], using the software OxCal ver. 4.3.2 [49–51]. All calibrated dates are presented in calibrated years BP with 95.4% probability range.

### Thermoluminescence dating

Additionally, the bottom-most layer of silty loams was dated with the use of thermoluminescence (TL) dating in the Department of Geomorphology and Quaternary Geology of the University of Gdańsk. The deposit moisture was measured in each sample. After drying, the dose rate (dr) was determined with the use of the MAZAR gamma spectrometer. The concentrations of ^226^Ra, ^228^Th, ^40^K (Table 2) in each sample were obtained from twenty measurements lasting 2000 s each. Equivalent dose (ED) was established on the 63e80 mm polymineral fraction, after 10% HCl and 30% H2O2 washing and UV optical treatment. The samples were irradiated with 20 Gy, 30 Gy, 40 Gy, 50 Gy and 100 Gy, dozes from ^60^Co gamma source. Before measurement, the samples were heated at 140°C for 3 hours. A sample pre-treated in this way was used to determine the equivalent dose (ED) (Table 2) by the TL multiple-aliquot regenerative technique [52], according to the description published by Fedorowicz et al. [53]. The registration of curves was performed on RA’94 (Mikrolab) thermoluminescence reader, coupled with EMI 9789 QA photomultiplier. The TL age was calculated according to Frechen [54]. A detailed description of the preparation and the equipment used in the study is contained in the paper by Fedorowicz et al. [53].

**Table 2.**
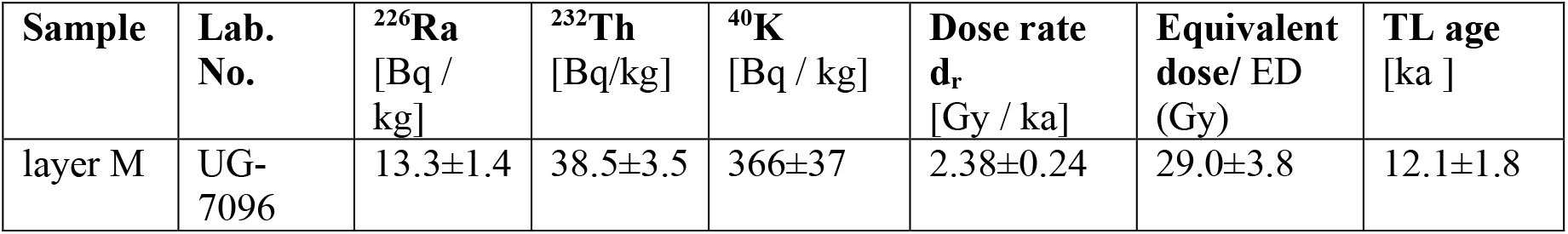
Thermoluminescence dating of a soil sample from Koziarnia, layer M.

| Sample | Lab. No. | <sup>226</sup>Ra<br>[Bq / kg] | <sup>232</sup>Th<br>[Bq/kg] | <sup>40</sup>K<br>[Bq / kg] | Dose rate<br>d<sub>r</sub><br>[Gy / ka] | Equivalent dose/ ED<br>(Gy) | TL age<br>[ka ] |
| --- | --- | --- | --- | --- | --- | --- | --- |
| layer M | UG-7096 | 13.3±1.4 | 38.5±3.5 | 366±37 | 2.38±0.24 | 29.0±3.8 | 12.1±1.8 |

### U-series dating

Due to the presence of radiocarbon dates reaching the limit of the method range, the U/Th method was additionally applied (Table 3). The chemical procedure was done in the U-series Laboratory of the Institute of Geological Sciences, Polish Academy of Sciences (Warsaw, Poland). The method included the thermal decomposition of organic matter and adding the ^233^U-^236^U-^229^Th spike to the samples, which were then dissolved in nitric acid. Uranium and thorium were separated from the hydroxyapatite matrix using the chromatographic method with TRU-resin [55]. The isotopic composition of U and Th was measured in the Institute of Geology of the Czech Academy of Sciences, v. v. i. (Prague, Czech Republic), with a double-focusing sector-field ICP mass analyzer (Element 2, Thermo Finngan MAT). The instrument was operated at low mass resolution (m/Δm ≥ 300). The measurement results were corrected to include background and chemical blank in the calculations. The age errors were calculated considering all uncertainties, except decay constant, using error propagation rules. Two cave bear teeth were dated and cross-checked with the use of radiocarbon method. All cave bear remains were assigned to *Ursus ingressus* according to preliminary analyses of ancient DNA as well as previous studies, which indicated that only this species (not *U. spelaeus*) was present on the territory of Poland in the Pleistocene [56–59].

**Table 3.** Uranium datings of Ursus ingressus teeth samples from Koziarnia. The reported errors are 2 standard deviations.

| Sample | layer | Lab. no. | U cont. [ppm] | U-234/U-238 | Th-230/U-234 | Th-230/Th-232 | Age [ka] |
| --- | --- | --- | --- | --- | --- | --- | --- |
| B 129 | F | 1124 | 0.0331±0.0002 | 1.14±0.01 |  |  |  |
| B 142 | G | 1125 | 0.0056±0.0001 | 1.19±0.01 |  |  |  |
| B 152 | G | 1126 | 0.0275±0.0001 | 1.22±0.06 | 0.200±0.009 | 86±4 | 24.3±1.3 |
| B 200* | I | 1127 | 0.0458±0.0002 | 1.16±0.06 | 0.209±0.007 | 55±2 | 25.6±1.0 |
| B 247 | L | 1128 | 0.0407±0.0002 | 1.27±0.08 |  |  |  |

\*Sample dated also with $^{14}\text{C}$ method (Table 1).

### Techno-typological analyses

The archaeological assemblage was analysed with the use of a geometric, morphometric and technological approach. A set consisting of 33 features was determined for each of the pieces of debitage. The attributes were divided into four general groups:

- general artefact morphology (the size, shape, state of preservation/fragmentation, symmetry, cross-section, profile, the character of the distal part),
- the condition of the dorsal side (the direction of the scars, cortex, interscar ridges, erasing chips, retouch),
- the condition of the ventral side (the bulbs, bulbar scars),
- the condition of the butt (the size, shape, profile, preparation).

In order to determine the characteristic features of the Jerzmanowician debitage, experimental studies were conducted additionally. They aimed at reproducing the bifacially-worked leafpoints and studying them to determine whether one can distinguish any specific features indicating leafpoint-shaping based on the morphology of the debitage. During the experimental session, two blades and two flakes were shaped by experimental knapper Miguel Biard into leafpoints, and the geometric morphometric features of the debitage were analysed.

### Traseology

Flint artefacts designated for traseological analysis were subjected to a cleaning procedure involving the use of warm water and acetone. The flint material was analysed using a Nikon LV150 metallographic microscope and a Keyence VH-Z100R digital microscope. The microscopic analyses were conducted using with a 50x to 400x magnification ratio. The noted macroscopic and microscopic traces – chipping, linear wear patterns and signs of usewear, linked to changes in the surfaces caused by post-depositional and utility factors. The listed traces were interpreted based on a comparison with an experimental reference database, kept together with the relevant documentation at the Institute of Archaeology of the University of Warsaw, as well as with reference to the appropriate literature.

The surfaces of the bone items selected for traseological analyses were cleaned using acetone. The optical-stereoscopic Olympus SZX9 and metallographic Nicon Eclipse LV 100 microscopes were used for the observation of the traces, along with the Nicon Shuttlepix digital microscope with a 6.3x to 100x magnification ratio. Natural and technological traces, as well as evidence of usewear were noted on the analysed material.

### ZooMS

The ZooMS (peptide mass fingerprinting) analysis followed protocols detailed elsewhere [60–62]. Both bone tools (KZ-2661.4 and KZ-2661.5) were sampled destructively (between 10-30 mg) and each bone sample was demineralised in 250 μl 0.5 M hydrochloric acid (HCL) at 4°C for 20 h. The samples were then centrifuged for 1 min at 10k rpm and the supernatant was removed. The demineralized collagen was then rinsed three times in 200 μl of 50 mM AmBic (ammonium bicarbonate) and 100 μl of 50 mM Ambic was added to each sample. Next, the samples were incubated at 65°C for 1 h. Afterwards, 50 μl of the resulting supernatant was digested with trypsin (Promega) at 37°C overnight, acidified using 1 μl of 20% TFA, and cleaned with C18 ZipTips (Thermo Scientific).

Digested peptides were spotted in triplicate on a MALDI Bruker plate with the addition of an α-Cyano-4-hydroxycinnamic acid matrix. MALDI-TOF-MS analysis was conducted at the Fraunhofer IZI in Leipzig (Germany), using an autoflex speed LRF MALDI-TOF (Bruker) in reflector mode, positive polarity, matrix suppression of 590 Da and collected in the mass-to-range 700-3500 m/z.

Triplicates were merged for each sample and taxonomic identifications were made through peptide marker mass identification in comparison to a database of peptide marker series (A-G) for all medium-to-larger sized mammalian species [62–64].

## Results

### Stratigraphy

The new fieldwork confirmed the massive destruction of the top part of the original sedimentary sequence. The topmost layer in the recent cross section, which can undoubtedly be correlated with the previous fieldwork, is layer K. It can be correlated with Chmielewski’s layer 12. There is no possibility to correlate the overlying strata J, B and A, as long as their remnants can be seen only close to the cave walls. All the overlying layers have already been destroyed, mostly due to 19^th^-century cave sediment exploitation.

In general, the sedimentary sequence can be divided into four lithostratigraphic series. The lowermost series is red residual clay of weathering origin (P), which has been locally preserved, a remnant of the older sedimentary series (S1). It fills the cracks and fissures in the bedrock. The second series consists of silt and sand (layer M), filling the bottom erosional rill, most probably a vadose canyon. The third series is built of red-brown and brownish loams (layers H’, I, H, I’ and L), containing highly corroded and rounded limestone clasts (Fig 3). The upper series consists of a set of grey loams containing either corroded or sharp-edged limestone clasts. It is divided into two parts by a lamina of red-brown clay (layer C). Layer K’, due to considerable amounts of charcoal, was a very dark black colour in Chmielewski’s trenches. In our trench, 40 m from the entrance, this layer still contains large charcoal fragments but it is more yellowish-dark grey in colour. The uppermost part of the section, especially strata situated above layer K’, has been disturbed. Layers younger than layer K are preserved only as remnants attached to the wall.

**Fig 3.**
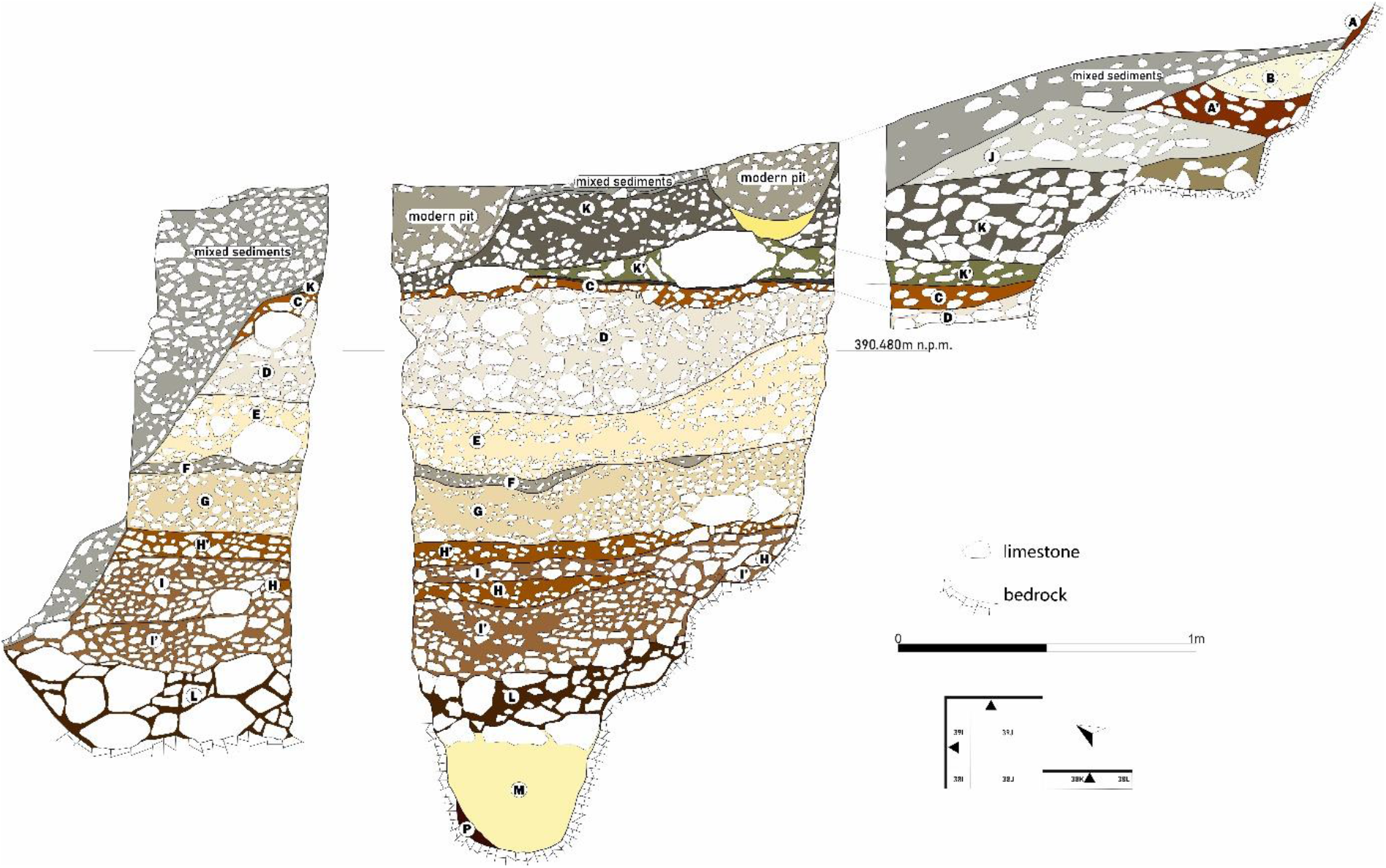
Northern and eastern cross-section of trench IX/2017 excavated in 2017 and located in the corner of the trench IX by W. Chmielewski (drawn by K. Skiba).

Several erosional surfaces can be identified within the sequence. The most prominent are situated at the bottom of layers I, E and D. They are marked as non-conformities. The most distinct is the bottom of layer I, which form erosional channels cutting into at least two of the lower layers H and I’.

### Chronology

In total, 14 animal bones and 9 charcoal fragments from the new excavations have been dated (Table 1). Radiocarbon dating was conducted on cave bear bones from the old Chmielewski excavations. Additionally, the date was estimated for two bone tools and the single cave bear bone from Römer’s collection (Fig 4).

**Fig 4.**
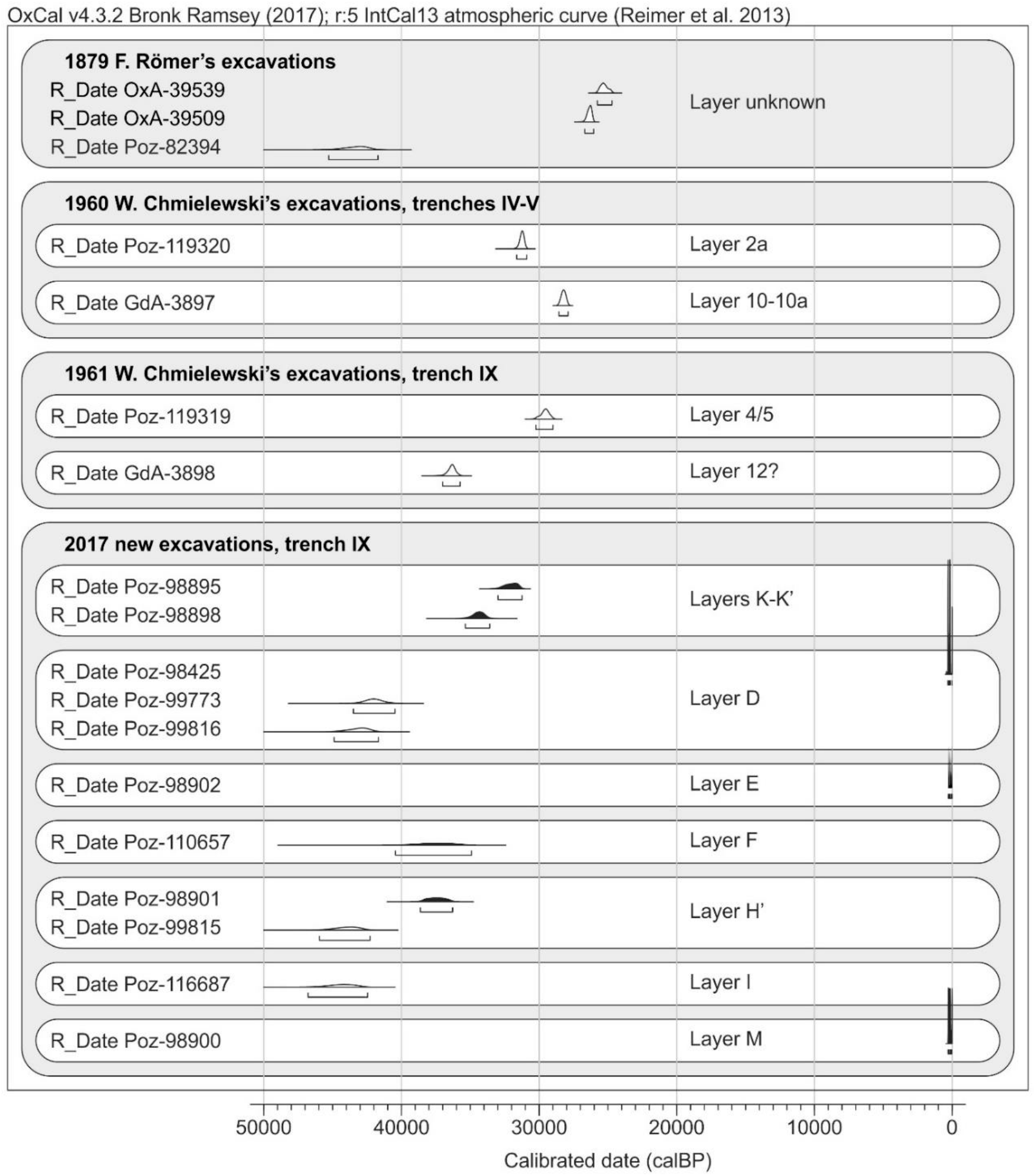
Calibrated radiocarbon dates from Koziarnia Cave arranged by layers. In black – the dates established for charcoal fragments, in white – for bones or teeth. The recent date Poz-98899, which falls beyond the IntCal13 calibration curve, is not shown, similarly to the open date Poz-99814.

The results show that at least some of the strata were contaminated by recent material. Recent dates were obtained solely for the charcoal fragments. This may indicate post-depositional processes connected with the extended exposition of the old open trench walls to external conditions.

Two bones were dated with the use of U-series (Table 3). In order to check the results, one of the bone specimens was dated with both the radiocarbon and U-series method. The U-series date is distant from the radiocarbon one (Table 1). This suggests that U-series dating is probably unreliable at this site. The reason behind this might be the open uranium system, i.e., the constant availability of uranium ions in the ambient sediments, which resulted in the continuous uptake of U from the environment by bone. In such cases, the U-series dates have only a *terminus ante quem* significance.

Among the newly established dates for the bones, 10 exhibit the atomic C:N ratio in extracted collagen which stays within the accepted range of 2.9-3.6 [65, 66] and indicates well-preserved collagen. One sample (date Poz-99806) yielded too low amount of collagen to measure the C:N ratio, while another (date GdA-3896) exhibited too low C:N ratio; therefore, we decided to discard these dates (Table 1).

The lowermost layers L-M yielded two radiocarbon dates. One of them represents recent contamination, the other was established based on material coming from the 1960s excavation; thus, we are not certain about its provenience. On the basis of the dating of the upper layers, layers L-M should be regarded as older than ca. 47 ky cal. BP. The TL date for layer M is incompatible with radiocarbon dating. This date could be biased due to the close proximity of bedrock, which resulted in a different radiation dose actually absorbed by the sediment than the dose assumed in the laboratory from the measurement of the concentration of radionuclides within the sample.

The complex of layers H’/I/H/I’ (H’ and I, including also the undated layers H and I’) exhibits a radiocarbon age of around 47-36 ky cal. BP. The overlying layer F (layer G was not dated) show a slightly younger age, of around 40-35 ky cal. BP. This chronology is based only on one date with a large margin of error (1200 years for 1σ, almost 2500 years for 2σ). The upper part of the sequence yielded dates which were not in chronological order with the lower ones. Material from layer D is as old as the material from the layer F and complexes of layers H’/I/H/I’. This indicates the erosion of material from the lower stratigraphic position and its re-deposition into layer D. The huge channel-like structure visible in the bottom of layer D supports this hypothesis. Layers K and K’ provided dates of around 35-31 ky cal. BP. This remains in accordance with the dating of layer F, especially if we consider the large margin of error for a single date from these layers. This also indicates that the erosion event followed by the re-deposition of layer D should be dated to around 37 ky cal. BP.

### Archaeological assemblage

During new fieldwork, over 1000 stone artefacts were collected. Table 4 shows the composition of artefacts found in each geological stratum. As a result of the opening of a new trench in the corner of the old collapsed trench and excavating not only *in situ* sediments, but also partly collapsed and moved layers, not all artefacts could be undoubtedly attributed to one level (Table 4). Within the majority of layers, the number of collected lithics has not exceeded 20 pieces each. Excluding mixed materials, only layers D, H’ and I’ are relatively richer.

**Table 4.**
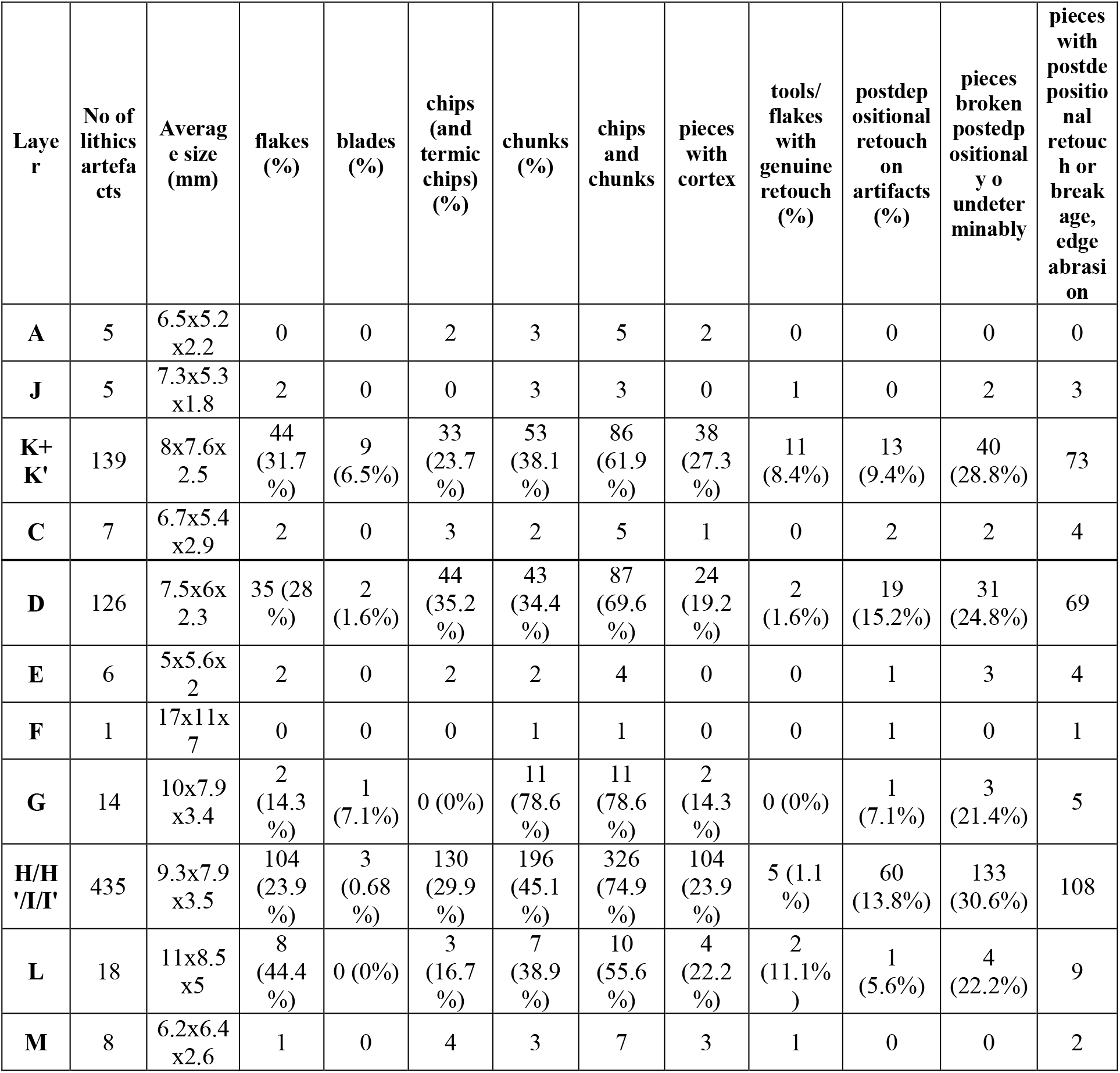
General composition of the Koziarnia Cave assemblage.

Most of the artefacts are tiny chips and chunks; on average, these constitute 69% of the assemblage. The average stone artefact size is 7.36 mm in length, 5.85 mm in width, and 2.65 mm thick. No cores or preforms were found within the assemblage. The small size of the artefacts can be explained by the location of the trench, which was situated 40 m from the entrance to the cave. One can expect scarce human activities to have been held so far from the entrance and the only source of sunlight.

Overall, 27% of all the flint artefacts had post-depositional retouches, and 35% of them had undergone post-depositional breakage. The highest impact of post-depositional processes was observed in layers I and I’. The state of preservation and small dimensions of the artefacts had a significant influence on further analyses. These factors made it significantly more difficult to identify any potential traces formed as a result of the use of individual flint specimens. The surfaces had been deformed to a high extent and covered with shiny or white patina. In addition, some parts of the edges had been destroyed, an effect of which were numerous post-depositional chippings, which stand out due to the distinctive “freshness” of the flake negatives as compared to the state of preservation of the remaining parts of the specimen surfaces.

Apart from layers K/K’, where several characteristic retouched pieces attributed to Gravettian were found, the other layers contained only uncharacteristic debitage, prevalently chips. To determine in which layer one could expect the Jerzmanowician occupation, analysis of the chips was required.

The general assumption was that in the Jerzmanowician assemblage, due to the characteristic retouching of the blades to shape a leafpoint, one could expect specific debitage, derived at the stage of the ventral thinning. Unlike in Middle Palaeolithic and Gravettian, we assumed that in the Jerzmanowician assemblage, one would be able to find chips and flakes with remnants of the ventral surface of the original blank/blade.

To test this assumption, we conducted experimental knapping and analysed the debitage obtained during the blade leafpoint shaping (Fig 5).

**Fig 5.**
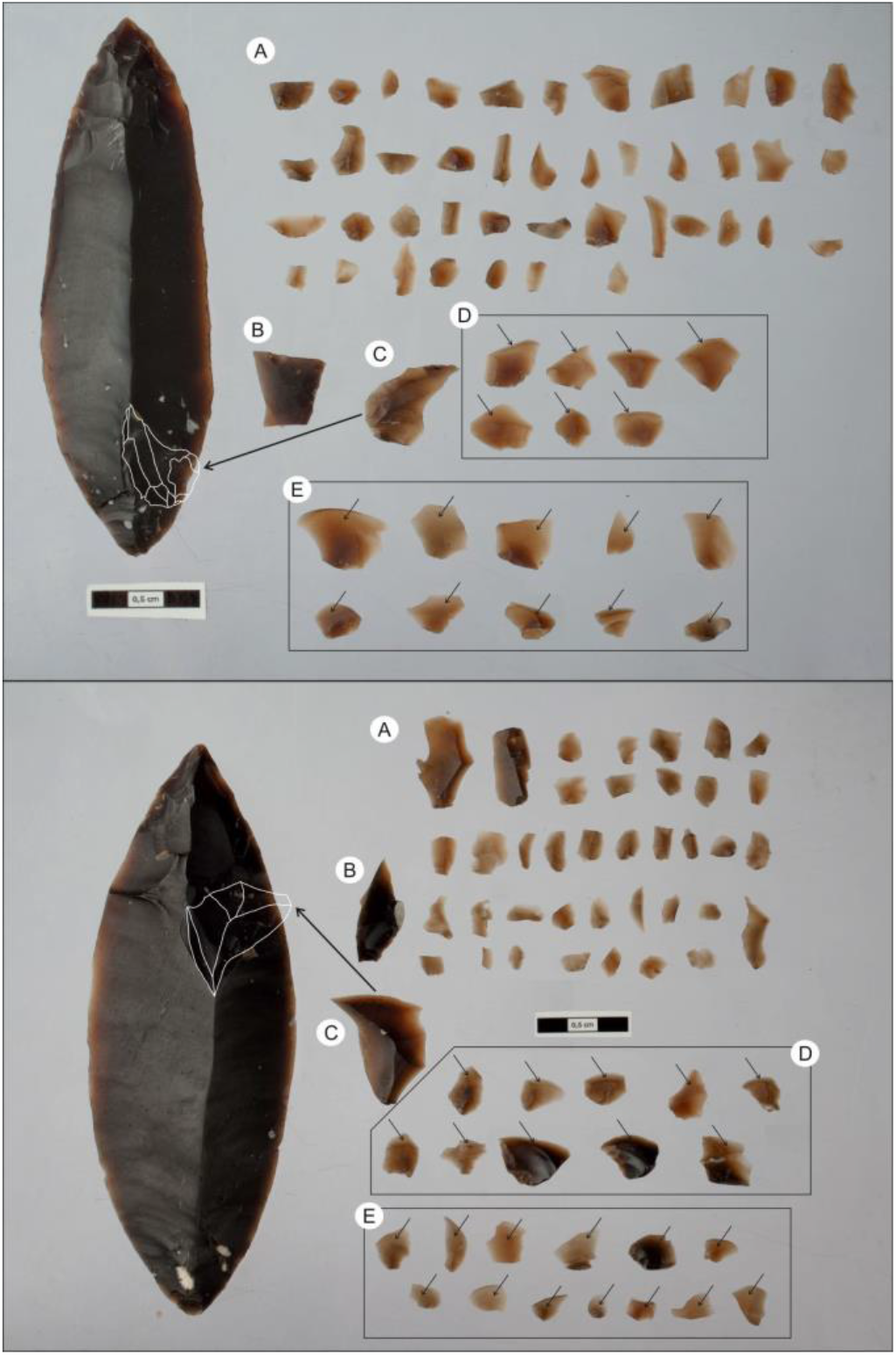
Results of experimental knapping, Jerzmanowician point and selected debitage products, (A) chips made during leafpoint production with ordinary morphology. (B) chip or flake detached during ventral thinning, near the leafpoint base. (C) chip/ flake curved due to reaching the transversal interridge of the original blank in its distal part. (D) The second generation of ventral thinning chips with remnants of ventral surface of the blank in their distal parts. (E) The first generation of ventral thinning chips with the ventral surface of the blank on their entire dorsal side.

### Small debitage: experiments

Experimental studies show that during the shaping of a blade into a leafpoint, approximately 150 chips over 2 mm in length are produced. Only several pieces had dimensions exceeding 1.5 cm and could be ascribed as small flakes. Chips are prevalent among the debitage. The chips can be divided into three general morphological groups.

The first group consists of ventral thinning chips (Fig 5E). They contain the remnants of the ventral surface of the blank on their dorsal side. They come from the initial thinning of the ventral side of the blade and can be described as the first generation of ventral thinning.

The second group contains chips which are slightly bent, with a width/length index of >1. On their dorsal side, they contain scars of even smaller removals in their proximal part and a big ventral scar on their distal part. Such chips come from second generation of ventral thinning (Fig 5D). Unfortunately, due to the small sizes of the analysed artefacts, in many cases, it is not possible to say if the remnant of the flat surface in the distal part of the artefacts is the ventral or dorsal side of the original blank.

The third group of chips consists of undeterminable uncharacteristic small chips (Fig 5A). They mostly come from dorsal thinning and shaping, as well as secondary ventral thinning. The characteristic feature in shaping a leafpoint out of the wide and rather thick blade is the presence of dorsal thinning chips, which reach the interscar ridge or the blank and contain remnants of such a ridge running transversally to their main axis (Fig 5C). Such flakes might be considered the specific debitage of leafpoint shaping.

Additionally, during ventral thinning near the butt part of the blade, in some cases a bigger chunk is produced, which aims to prepare the correct angle for further removals (Fig 5B).

Based on experimental studies, one can assume that only the presence of chips from the first generation of ventral thinning can be treated as evidence for ventral thinning, and - therefore - can be associated with leafpoint shaping. Although the presence of only the second generation’s chips cannot be indication for leafpoint production, their appearance together with first generation chips could provide additional support for such an assumption.

Table 5 presents morphological analysis of the chips found in distinct strata in Koziarnia. The results show that ventral thinning chips could only be found in layer D (Fig 6) and in the disturbed sediments (n=28). In other layers, the undeterminable chips of the third type prevail. The presence of ventral thinning chips leads to the assumption that layer D should be associated with an assemblage that used a ventral thinning method, probably bifacial leafpoint shaping.

**Fig 6.**
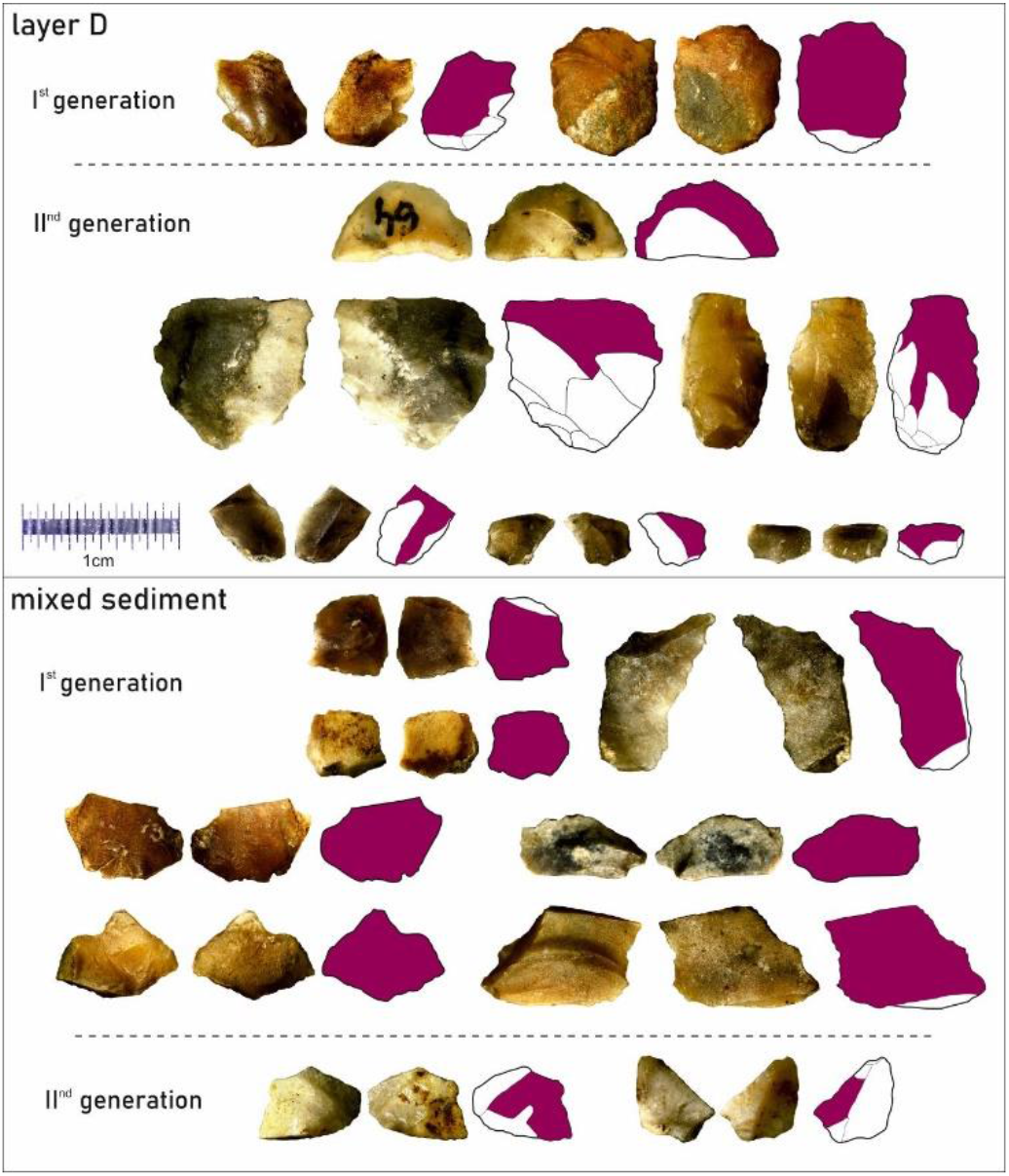
The first of second generation of ventral thinning chips containing remnants of the ventral surface of the blank (marked in pink) on their dorsal side. Artefacts found in layer D and in the mixed sediment, trench IX/2017 in Koziarnia Cave.

**Table 5.** Comparison of features determined in chips obtained during experimental leafpoint shaping, with the archaeological material from layers K/K’, D and H’/I/H/I’/L indicating the presence of ventral thinning chips in layer D.

| Feature/Layer | Experimental assemblage | Layers K/K' | Layer D | Layers H'/I/H/I' and L |
| --- | --- | --- | --- | --- |
| Total number of chips |  | 32 | 44 | 133 |
| *First generation of ventral thinning [flake/chip with double ventral surface]* | ++ | - | ++ | - |
| *Second generation of ventral thinning [chips with remnants of ventral surface]* | ++ | - | ++ | - |
| *Second generation of thinning [bent short chips]* | ++ | (+) | ++ | ++ |
| *Flake/chip with orthogonal scars* | ++ | (+) | + | - |
| *Curving flakes/chips from dorsal thinning* | + | - | - | - |
| *Thin elongated chips* | ++ | - | ++ | (+) |
| *Presence of lip* | ++ | + | + | ++ |

### Other artefacts

#### Holocene

The upper parts of the mixed sediment provided a minimal number of Holocene period finds, consisting of pottery, a glass artefact and metal objects. The ceramic assemblage is highly fragmented and poorly preserved. It comprises 6 pieces of uncharacteristic prehistoric pottery sherds, 1 fragment of Roman-period wheel-turned ware fired in reducing atmosphere, and 2 pieces of vessels dated tothe 18^th^ or 19^th^ century, made of white clay and covered with yellow glaze. The find assemblage is supplemented with a small fragment of a patinated glass artefact, possibly a vessel, and small pieces of undefined metal objects. Due to their poor preservation, the chronology of these finds must remain uncertain.

Considering the recently discovered Holocene period finds, it is fair to say that they do not bring new data to the studies of the use of Koziarnia in late prehistory and historical times. Some more detailed insights into this topic were provided thanks to previous research campaigns, which were focused on the entrance zone to the cave. As evidenced then, the site was extensively used since the Neolithic up until the modern period [27]. The small amount of Holocene period finds from the 2017 excavations has to be seen in the context of the distance of the trench from the cave mouth.

#### Layer K, K’

Both horizons are described together since layer K’ has the same petrological features as layer K. The only difference is the presence of a high concentration of charcoal in layer K’, which changed the colouration of the layer. Therefore, one can assume that layer K’ is a human occupation episode within the accumulation of layer K. Due to the significant destruction of the top levels, layers K and K’ were visible only in a tiny area of ca. 1 m^2^. The lithic assemblage consists of 139 artefacts. It should be noted that materials from the layer determined ad mixed + K were also included.

Interestingly, almost half of the artefacts from layers K and K’ have traces of fire, which goes well with the high concentration of charcoal. The majority of artefacts were post-depositionaly damaged. Out of the 53 flakes and blades, only five were found unbroken. Nonetheless, the assemblage contains clear Middle Upper Palaeolithic i.e. Gravettian, elements represented by five fragments of backed bladelets (Fig 7 FLO/A34/17, FLO/A45/17a, FLO/A96/17, FLO/A45/17c, FLO/A45/17e), an endscrapper (Fig 7 FLO/A45/17d) and a double microburin, reworked into a double perforator (?) (Fig 7 FLO/A45/17b). Besides the tools mentioned above, one burin spall has been noted, as well as a small bladelet, which could be either a burin spall or crested blade. The assemblage consists of blades and bladelets and is distinct from the other assemblages due to the use of excellent quality almost translucent Jurassic flint raw material. However, not much can be said about the technology and morphometric characteristics of the assemblage, as mostly medial and distal flake and blade fragments were recovered.

Use-wear traces - linear traces and impact fractures [67] have been observed on four small backed bladelets, based on which it could be assumed that they were used during activities linked to hunting (Fig 7: FLO/A34/17, FLO/A45/17a, FLO/A96/17, FLO/A45/17c).

The first of them (FLO/A34/17) is characterised by the lack of any distinct post-depositional traces. In its middle fragment, linear traces were observed running outward from the chipping negatives. These marks are located on the lateral edge, intentionally left unretouched. The distinguished linear traces are located parallel to the tool’s axis of symmetry. The placement of these marks indirectly indicates the method of depositing it, i.e. with the backed bladelet parallel to the shafts. Additionally, the breakage of the tip was observed to have a straight profile, which could be associated with hunting weapons, but fracturing of this kind is not distinctive for this type of activities.

In turn, the second backed bladelet (FLO/A96/17) should be noted for its characteristic breakage of the tip. The impact fracture (hinge terminating bending fracture) has been identified. This type of breakage morphology usually enables to link the tool with hunting weaponry. In addition, the linear traces observed in the middle part of the tool can be connected, due to their underdeveloped form, with its use as a projectile, but simultaneously the influence of post-depositional factors cannot be excluded.

The third backed bladelet (FLO/A45/17a) is characterised by an impact fracture (step terminating bending fracture) in its bottom part. The macroscopic morphology of the trace suggests to a certain extent that the described backed bladelet might have been used as a hunting weapon.

It cannot be excluded that also the next specimen (FLO/A45/17c) was used during hunting. This is the upper fragment of a backed bladelet broken in a unique manner; one end of the breakage with a concave profile is elongated. The artefact might have been the tip of arrowhead projectile.

#### Layer D

Compared to other layers, this one contained a relatively rich assemblage (n=137), although the materials were heavily damaged, mostly through breakage (Table 4). Among the 90 blades, bladelets and flakes, only six were unbroken. Most of the flakes represent undeterminable debitage; however, at least some of them have negatives attesting to bidirectional knapping (Fig 6), which is a characteristic feature of Jerzmanowician [14, 21]. A single flake has the features of a bifacial shaping flake.

On the same level as layer D, but in the disturbed sediment of the old trenches, a blade with ventral thinning of the bulb was found (Fig 8: A1/17). The artefact may be interpreted as the broken part of a leafpoint; however, it contains numerous post-depositional retouches, which changed its shape. The usewear traces located along its longitudinal edges but due to the underdeveloped form of the polishes their detailed origin cannot be determined. However, the provenience of the usewear traces is unclear.

**Fig 8.**
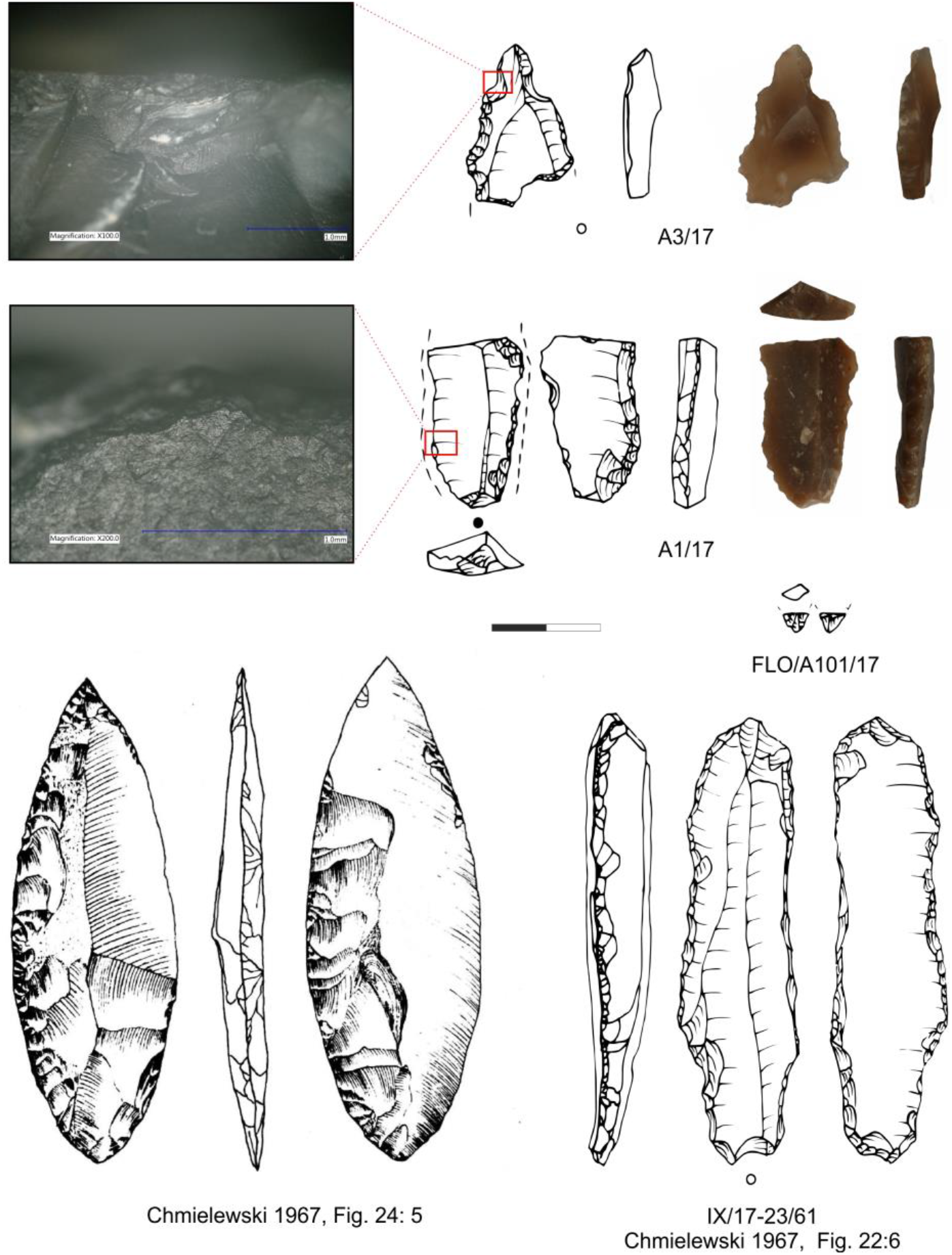
Artefacts from Koziarnia Cave attributed to Jerzmanowician. A3/17, FLO/A101/17 found in trench IX/2017 in layer D; A1/17 found in mixed sediment in trench IX/2017; Leafpoint was found by F.Römer in the second half of 19^th^ century [24, 27]; IX/17-23/61-blade made on double platform core found by W. Chmielewski in layer 17 of trench IX, later called layer 15 in the final publication. [27].

To conclude, one can see at least several features indicating that one is dealing with traces of Jerzmanowician occupation in layer D. The most prominent among these are the above-described presence of the ventral thinning chips (Fig 6) and the debitage with bidirectional scars. A bidirectional knapping scheme is confirmed also by a big blade detached from a bidirectional core found in the same layer by W. Chmielewski (Fig 8: IX/17-23/61).

#### Layers E, F, G

These three layers represent one geological sedimentary event affected by human occupation, which can be traced in layer F. This layer is characterised by a high concentration of charcoal, but only a single artefact was found inside (Table 4). The above and underlying layers E and G contained in total 21 artefacts (Table 4). They consist of uncharacteristic elements, with a single bifacial shaping flake from layer E. The cultural attribution of the assemblage is impossible.

#### Layer H, H’, I, I’

The assemblage found in the lowermost layers consists of 436 artefacts (Table 4). The flakes represent only undeterminable debitage. The edges are heavily damaged due to post-depositional retouches creating pseudo-retouched tool-like artefacts. The pseudo-retouches are present even on the 0.3-cm-long chips, indicating the intensity of the post-depositional damage. At least ten chips and flakes can be described as bifacial thinning debitage due to their knapping angle (60°-70°).

The assemblage contained one endscraper (Fig 9: A9/17), and possibly one “groszak” (Fig 9: A6/17). No usewear traces were found on these artefacts. A single flake contains a multiscarred butt in the shape of a *chapeau de gendarme*. All the described features indicate that these layers should be attributed to the Middle Palaeolithic. Unfortunately, the small size of the debitage and a high post-depositional damage unable more detailed cultural attributions.

**Fig 9.**
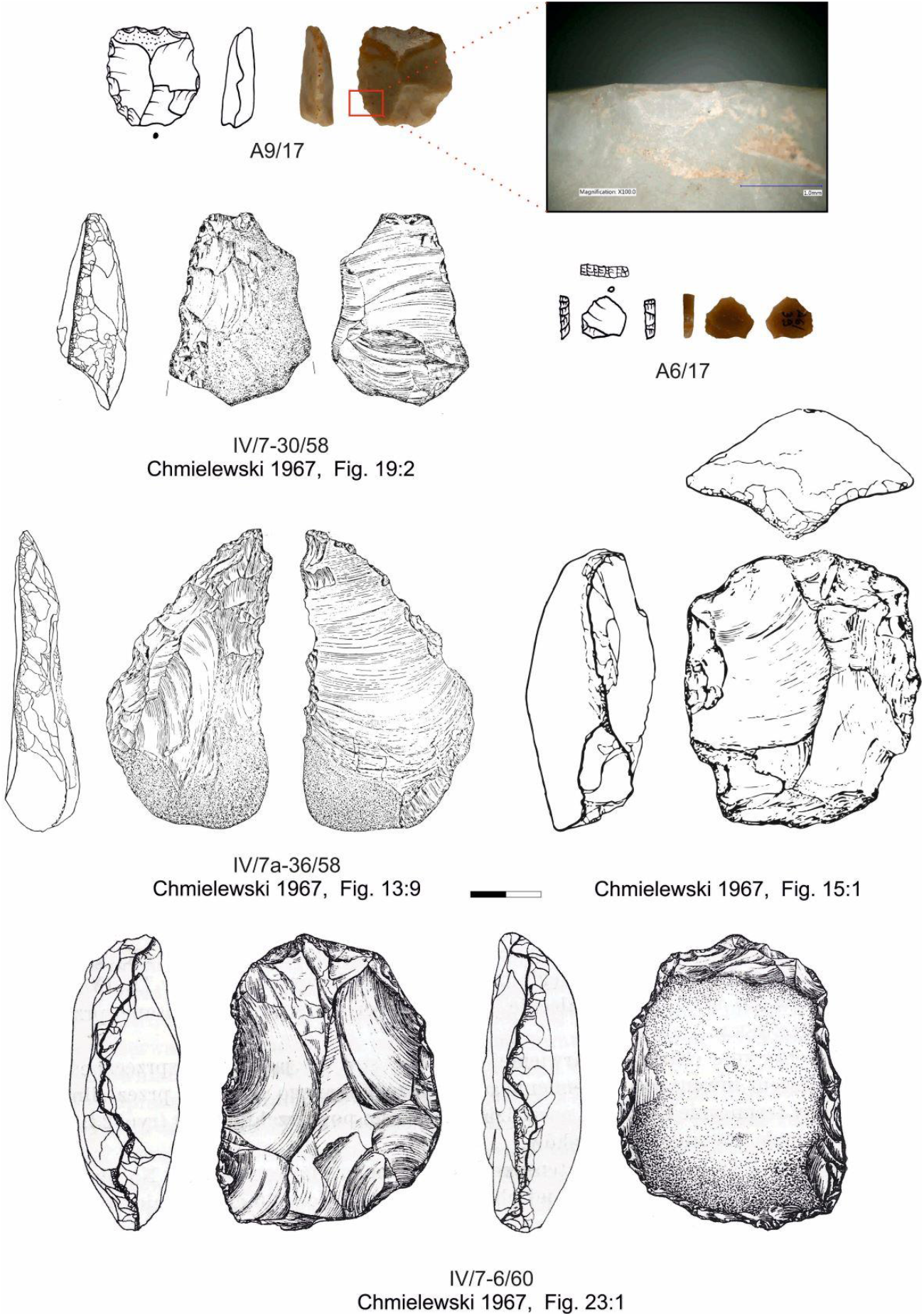
Middle Palaeolithic artefacts from Koziarnia Cave found recently in layers H’-L and by W.Chmielewski in layers 17-20 [27].

#### Layers L, M

They include 26 artefacts (Table 4) containing three retouched flakes and a single bifacial shaping chip (Table 5).

### Bone Artefacts

Two unpublished bone tools from Koziarnia cave were found in F. Römer’s collection. One of the tools is a short broken piece of bone with a smoothened ending (Fig 10:1). Numerous lengthwise cracks and chippings linked to exfoliation were observed on this artefact. One of the endings has been broken as a result of natural factors, while the other was formed diagonally through being burnished on a stone pad. No traces of usewear enabling the identification of its function were observed on the artefact.

**Fig 10.**
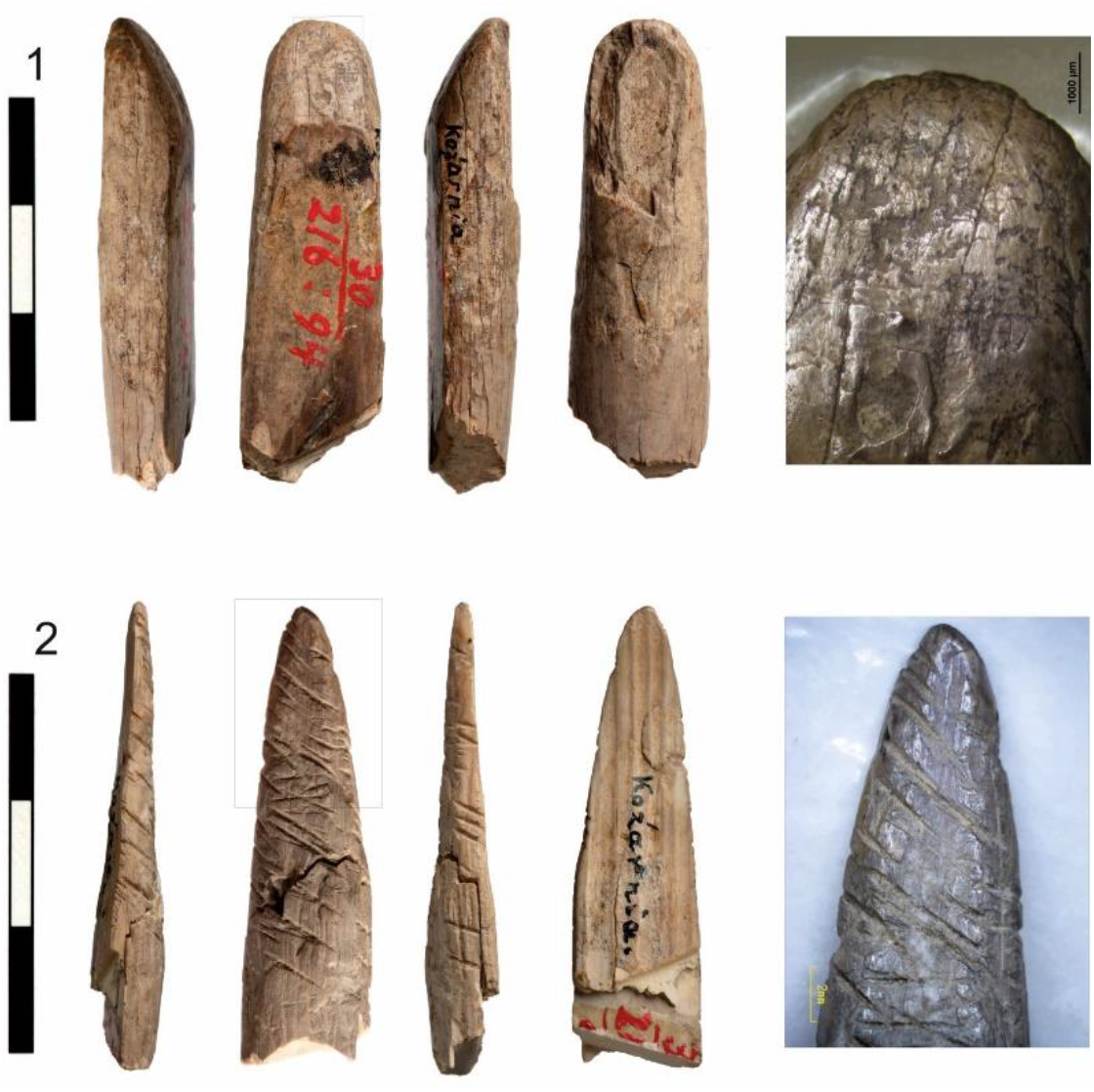
Bone artefacts found by F. Römer in Koziarnia Cave. (1) Fragment of broken bone with smothered ending. (2) Fragment of bone point with incisions. Photo M. Bogacki

The second piece is a part of a bone point with multiple incisions on its outer surface (Fig 10:2). Wide linear marks of different depths, overlapping each other and parallel to the longer axis of the artefact, were observed. They had been formed during the shaping of the blade through being scraped by a flint tool. On the entire surface of the blade, there are distinct, deep and wide diagonal notches located parallel to each other, only intersecting in the middle part of the tool. They were made with a flint flake or chip through repeated sawing backwards and forwards. This type of notch should be seen as a kind of artefact decoration. The surface of the artefact is smoothened and useworn, especially at its tip.

The spectra obtained from both bone specimens through ZooMS analysis have been taxonomically identified as Elephantidae. The marker series are similar for some closely related species. In this case, possible species can belong to the *Elephas, Mammuthus*, and *Palaeoloxodon* genera. Considering the archaeological context, these two bone tools were most likely manufactured from woolly mammoth remains.

Both artefacts obtained similar radiocarbon dates of 25-26 cal. ky BP (Table 1), which are younger than the chronology of the uppermost layers excavated in 2017. These radiocarbon dates indicate the presence of later Gravettian occupation in Koziarnia. It was most probably connected to one of the layers already destroyed in the cave.

## Discussion

### Koziarnia – correlation of layers

The stratigraphy observed in the 2017 trench fits well the description and documentation of the stratigraphy of trench IX by W. Chmielewski. The only difference is the presence of at least four separate strata, which were treated as a unified layer 17 by W. Chmielewski. Based on the new fieldwork, one can differentiate at least four substrata within the layer 17, differing due to the presence of weathered limestone clasts and the colouration (from the top: layers H’, I, H, I’). Traces of the relatively most intensive human occupation were found in the uppermost layer H’ and the lowermost layer I’.The second difference between the cross section presented by Chmielewski and the recent study is the relatively small amount of charcoal found in layer K’, which can be correlated with layer 13 by Chmielewski. Nonetheless, this layer contained the highest number of charcoal fragments in the entire sequence but their concentrations did not change the colouration of the stratum, as had been observed by Chmielewski in trenches IX and especially IV & V. As long as these layers can be correlated with human occupation, one can presume that the highest charcoal concentration could indicate an occupation zone, which weakens as it nears the end of the cave corridor.

The comparison of all the available drawings of the cross sections from all the previously conducted fieldwork enables to reconstruct the general correlation of the layers (Fig 11). Based on such correlation, one can see that the trenches located in the entrance zone revealed the presence of a thick Holocene sequence of humic horizons and underlying loess layers, which can be divided into two separate horizons. The loess sediments, through a comparison to other caves in the region, can be correlated with units ‘A’ and ‘C’, according to the lithostratigraphic scheme by Krajcarz et al. [68] and dated to late MIS 3 and MIS 2. However, we don’t have any direct dating data. All the older strata were probably washed away from the entrance zone before the late MIS 3. The most problematic issue linked to the whole stratigraphy of the Koziarnia Cave is the almost absolute destruction, removal and mixing of the sediments of the main chamber, which was probably the central settlement zone with the highest concentration of artefacts. The uppermost layers located inside the cave were almost completely removed in the 19^th^ century. The remnants of the original stratigraphy can be found only attached to the regolith visible on the wall of the main corridor. The current cave infilling only contains layers dated to MIS 3, except for the lowermost strata M (21) and P, which might be older.

**Fig 11.**
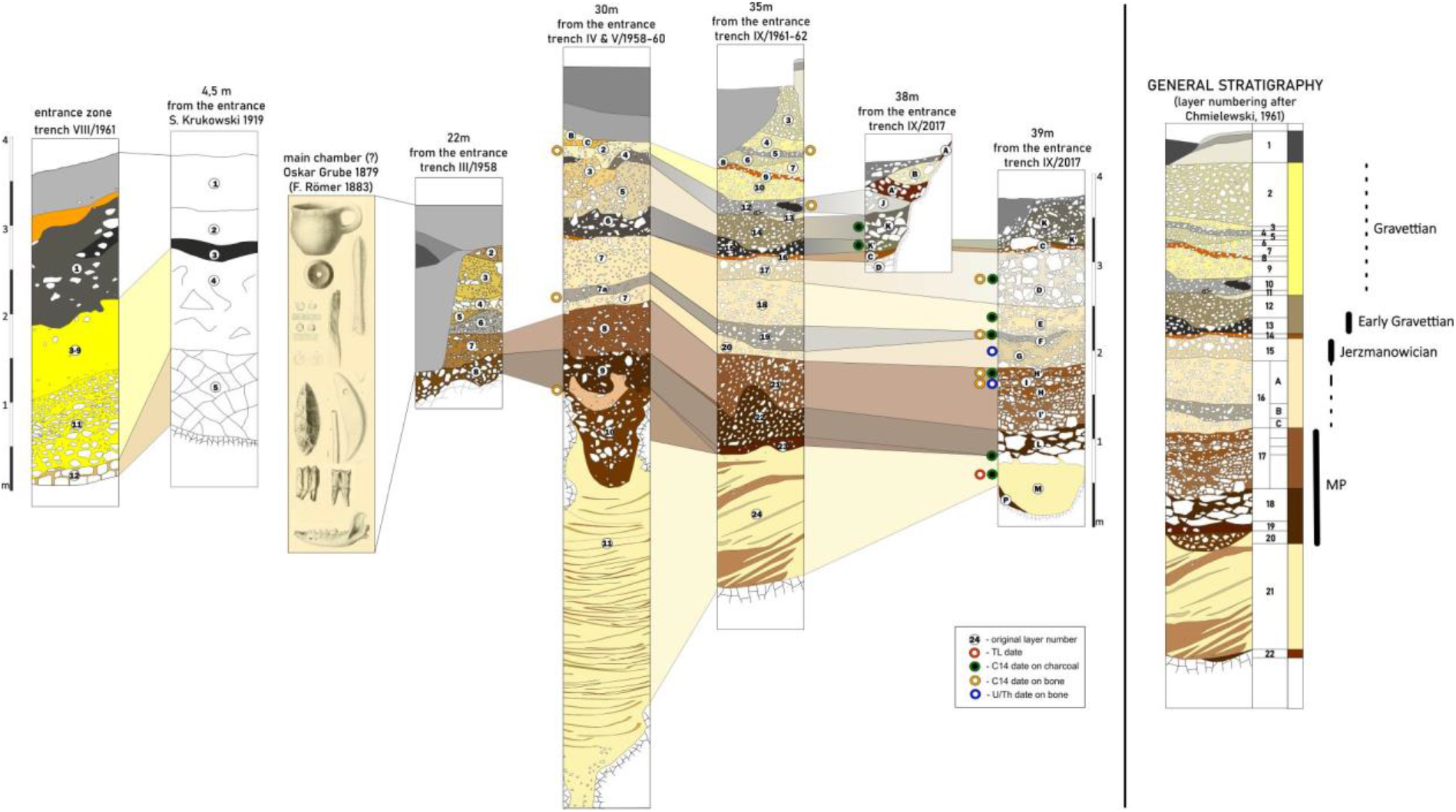
Correlation of all the profiles obtained during subsequent archaeological fieldworks in Koziarnia Cave with a location of samples used for dating. S. Krukowski field documentation [after 35, 36]; artefacts found by F. Römer [24]; W. Chmielewski profiles redrawn after Chmielewski [27]and a field documentation of trench VIII, trench IV, V & IX.

Based on the artefacts found and the obtained dates, one can see at least four different Palaeolithic settlement episodes in the cave. The Middle Palaeolithic is connected to layers 17(H’/I/H/I’) and 18 (L), Jerzmanowician – to layers 15-16 (D-E-F-G), and the Early Gravettian – to layers 13-12 (K-K’). The later Gravettian episode (25-26 ky BP) cannot be attributed to any particular stratum but was manifested by the presence of two bone tools found in F. Römer’s collection.

The general correlation of the layers shows that the amount of charcoal in the Early Gravettian horizon diminishes towards the end of the cave, and is the most intense approximately 20-25 m from the entrance. In contrast, the thickness of the Jerzmanowician layers 15-16 increases as they near the end of the cave (a 75-cm-thick layer is 40 m from the entrance), while they disappear towards the cave entrance.

### Chronology

Most of the established dates cover the period between 46 and 24 ky cal. BP. However, several charcoal datings provided unexpectedly recent ages (Table 1). Based on the taxonomical analysis of the charcoal assemblages, it can be assumed that a part of the floated samples were indeed contaminated and this can be confirmed by the presence of a few samples containing singular findings of charcoal fragments belonging to fir *Abies alba*, hornbeam *Carpinus betulus* and beech *Fagus sylvatica*, trees that are considered late-coming species in the vegetation history of Poland [69]. Such samples came mostly from areas located near the previous excavations. After preliminary analysis, the existence of post-depositional disturbances was confirmed, indicating these places as ones that should be excluded from any chronological inference. However, in a few other samples, only coniferous taxa were found (juniper *Juniperus communis* and pine *Pinus* type *sylvestris-mugo*), which suggests that they could have originated from Pleistocene layers, but their radiocarbon dating showed modern contamination (Table 1). This analysis has evidenced that the very meticulous study of strata in the context of all archaeological and biological findings is needed to understand taphonomic processes in cave sites.

Another explanation for the observed discrepancy between the dates achieved from bones and at least some dates achieved from charcoal, includes the altered ^14^C content in charcoal fragments. From recent study [70] we know that carbon in wood during the high temperature processing (such as burning) is a subject to kinetic fractionation of isotopes. Namely, charcoal from coniferous wood burned in low temperatures is enriched in heavy carbon and oppositely, burned in higher temperature (400-600 °C) is depleted in heavy carbon in relation to the original wood. If dated charcoals became from burning in relatively low temperatures, e.g. in the outer part of fireplace, this may likely produce a shift to younger radiocarbon dates.

It is worth noting that the radiocarbon dates from layer D are not in the correct order with those from the lower strata (Fig 4). This can be an effect of the mentioned isotopic fractionation in burnt wood, or likely the effect of re-depositional episodes, possibly related to the erosional structures visible in layers E and D. The directions of this transport are difficult to reconstruct as the 2017 excavation area was quite small and delivered minimum data on the geometry of sedimentary structures. Moreover, the topography of the cave floor was disturbed here due to the previous exploitation. However, the higher elevation of sediments in the area closer to the entrance (especially visible in W. Chmielewski’s trench at the 30th metre) may suggest that this area served as a source of material for colluvial activities. If we adopt the hypothesis that at least some dates from the upper layers represent a re-deposited material, we need to accept that the faunal, anthracological and archaeological assemblages from these layers could also have been affected by colluvial mixing.

If we look at the distribution of the probability density of radiocarbon dates regardless of the stratigraphy (Fig 12) we can detect several phases of the deposition of dated material. The phases of deposition of the animal remains took place ca. 46-41 ky cal. BP, ca. 37-35.5 ky cal. BP, and ca. 32-28.5 ky cal. BP. The dates from charcoal fragments are restricted to ca. 39-31 ky BP, while the dated bone tools to ca. 26.5-24 ky cal. BP. Assuming that charcoal fragments are the remnants of hearths, the probability density of radiocarbon dates for charcoal represents the human settlement phase. During this phase, we can identify three weakly separated subphases. The first one can be dated to ca. 39-36 ky cal. BP (represented by two dates) whereas the second to ca. 35-33.5 ky cal. BP (a single date), and the third to ca. 33-31 ky cal. BP (a single date). Due to the stratigraphic position of the samples, we may assume that the first phase is connected with Jerzmanowician occupation, while the second and third with Early Gravettian. The last human settlement phase in 26-24.5 ky cal. BP also represents traces of the Gravettian occupation. Another phase, not shown in Fig 12, is the modern one (around 300 y BP until modern times), based on the most recent dates achieved for the charcoal.

**Fig 12.**
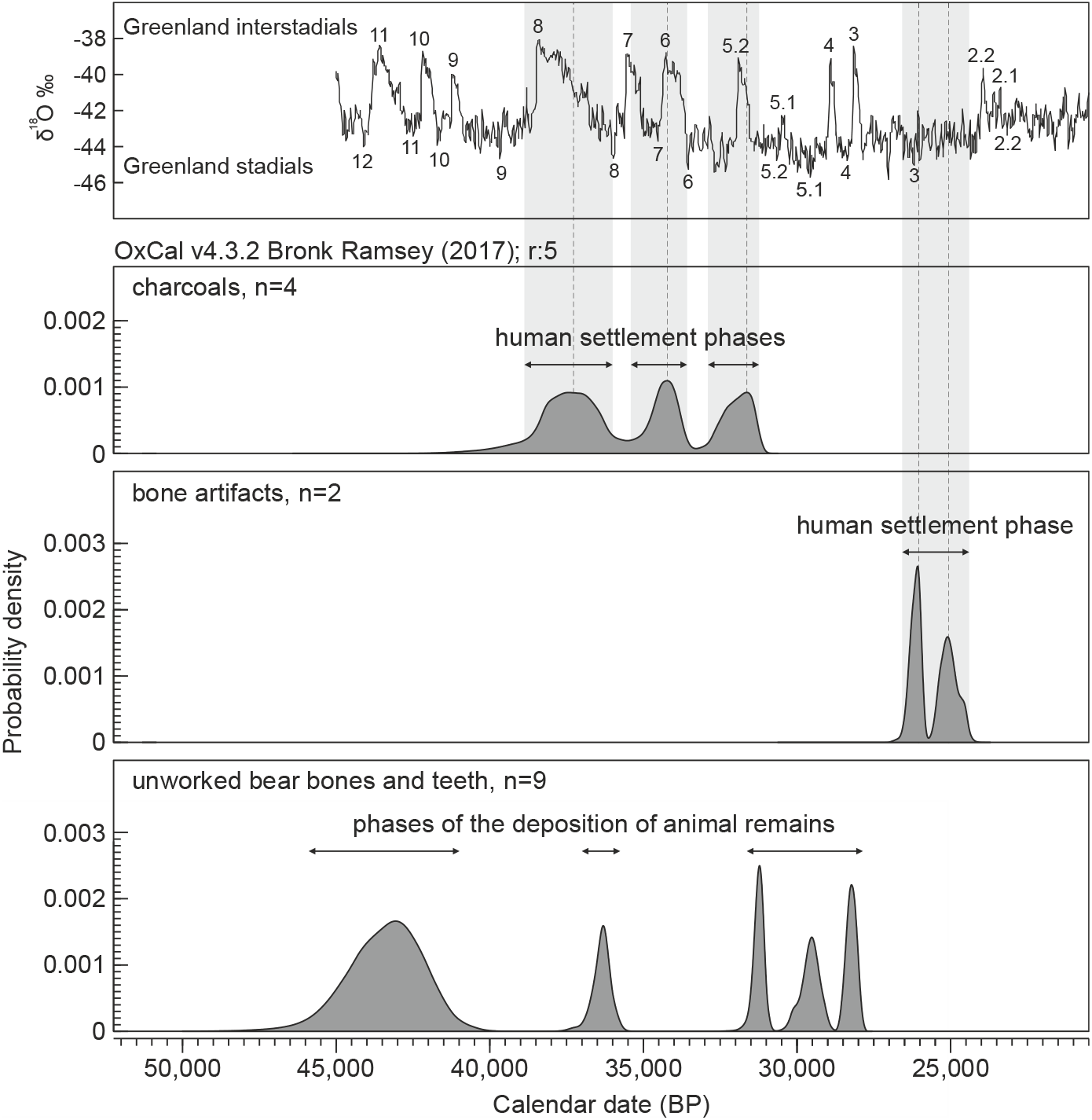
Distribution of the probability density of calibrated radiocarbon dates for Koziarnia Cave obtained for charcoal fragments (pink) and bone tools (yellow) compared with the revised δ18O curve in the Greenland ice core (blue) obtained by combining the Cariaco Basin (Hulu Cave) and Greenland ice core (GICC05) records [73]. Corresponding Greenland stadials (GS) and interstadials (GI), as determined by Rasmussen et al. [74] and Seierstad et al. [75] are indicated by numbers placed below or above the δ18O curve, respectively. The four most recent dates are excluded.

It is worth taking note of the alternate occurrence of dates determined for charcoal fragments and animal remains (Fig 12). All the dated animals were bears (mostly the cave bear, but one date has also been established for the brown bear). Bears used the caves as hibernation dens, and their presence in a cave could not be contemporaneous with human settlement [71, 72]. Our dataset indicates that Koziarnia Cave indeed had been alternately occupied by humans and bears.

In Fig 12, we compared the distribution of the probability density of radiocarbon dates with the pattern of the revised δ^18^O curve in the Greenland ice core. The curve represents reliable climate proxy reflecting global climatic changes in the Pleistocene [73–75]. If we take into account dates obtained for charcoal fragments and bone tools, which are direct indicators of human settlement in Koziarnia Cave, we can notice interesting relationships. Three peaks of the charcoal date distribution well correspond to three warmer periods (Greenland interstadials, GI-8, GI-6 and GI-5.2), whereas the minima of this distribution coincide with colder periods (Greenland stadials, GS-8 and GS-6). Although the date distribution for bone artifacts is shifted to Greenland stadial GS-3, there is a clear gap between these two distributions, which corresponds to the coldest stadial GS-5.1. This result suggests that Koziarnia Cave could be inhabited by human groups in waves especially in warmer periods, whereas climate cooling could discourage people from settling in this place.

### Other sites – correlation of profiles

Jerzmanowician assemblages are known from Nietoperzowa, Mamutowa, Puchacza Skała and Shelter above the Zegar Cave sites in Poland [20, 76–79]. The stratigraphic correlation of Jerzmanowician-bearing strata from Koziarnia Cave and Nietoperzowa Cave was studied by T. Madeyska-Niklewska and was first presented by Chmielewski et al. [27] and then by Madeyska-Niklewska [80]. According to this interpretation, layer 15 (D) from Koziarnia, where we found traces of Jerzmanowician occupation, correlates with layer 10b of Nietoperzowa Cave. However, the Jerzmanowician settlement is well-known from the younger layers 4-5-6 of Nietoperzowa Cave. It is difficult to compare the sequences from both caves based on the lithology, as they represent rather different facies (Fig 13). Sediments from Nietoperzowa Cave are mostly silty loams with limestone clasts, deposited in the near-entrance area under the strong influence of aeolian activity. In Koziarnia Cave, the recognized sediments are coarser, they were deposited quite deep inside the cave, and they are mostly limestone debris. However, the proportion of angular to subangular clasts, presented by Madeyska-Niklewska [80], enables correlating the Jerzmanowician layer 15 (D) from Koziarnia Cave with either the lowermost or the uppermost Jerzmanowician-bearing strata from Nietoperzowa Cave, namely layers 6 or 4.

**Fig 13.**
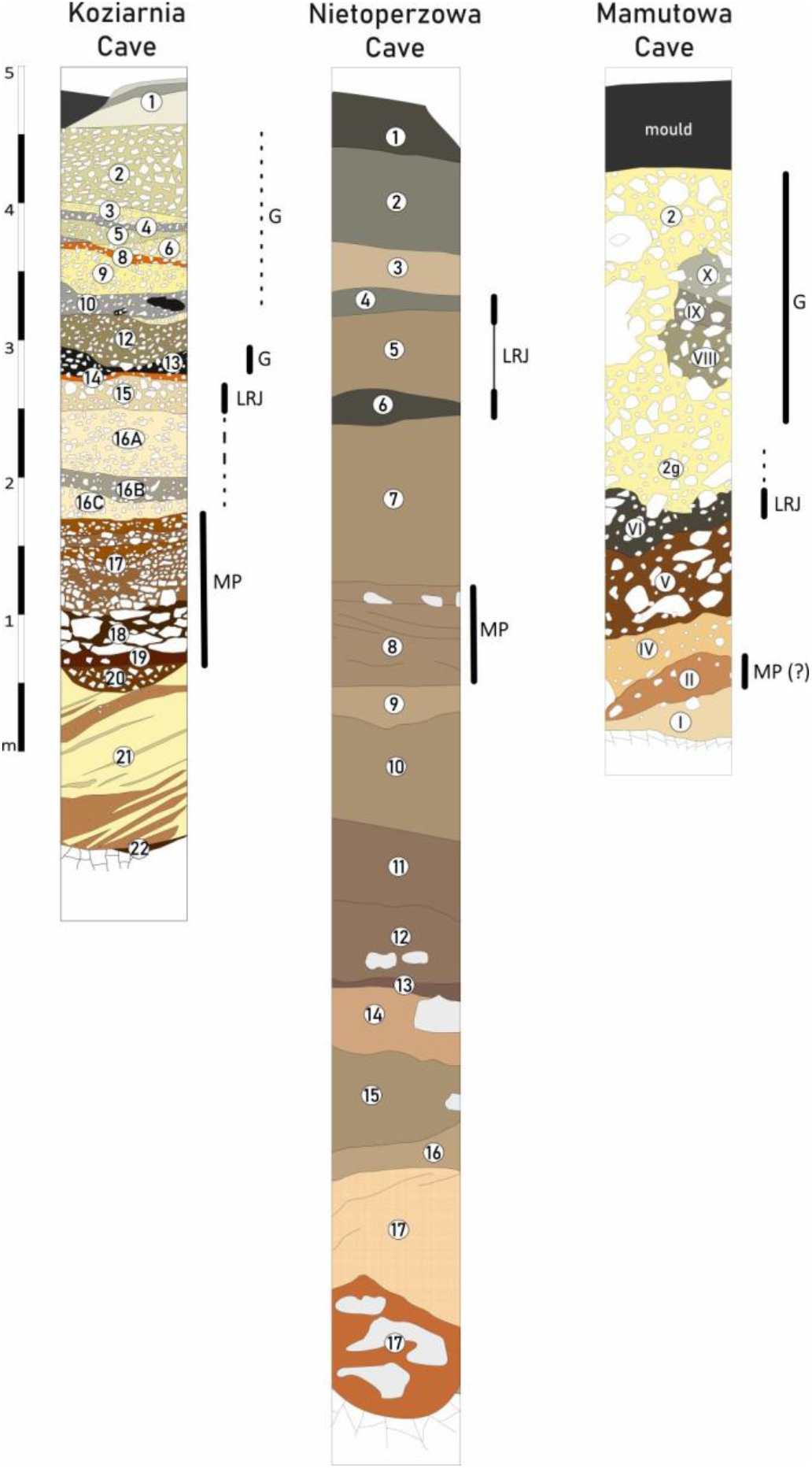
Correlation of profiles from three Jerzmanowician sites: Nietoperzowa Cave [33], Mamutowa Cave [81, 82] and Koziarnia Cave. Symbols used: G-Gravettian; LRJ - Lincombian-Ranisian-Jerzmanowician; MP - Middle Palaeolithic.

In Mamutowa Cave, the Jerzmanowician assemblage was found in layer VI [76, 81–83], which was blackish due to a significant concentration of charcoal (Fig13). Unfortunately, a detailed stratigraphic comparison between Koziarnia and Mamutowa caves are restricted due to the different character of the sediment at both sites. Kowalski’s layer V overlying the Jerzmanowician horizon was mostly loess sediment, which is not present inside the gallery in Koziarnia.

Interesting conclusions can still be derived from a comparison of the radiocarbon chronology of Koziarnia Cave with other sites of the Middle/Upper Palaeolithic transition and early Upper Palaeolithic from southern Poland (Fig 14). The chronology of the Jerzmanowician assemblages is currently based on 38 radiocarbon dates [S3, 33]. The majority of the radiocarbon datings were obtained on either cave bear (n=29) or bird (n=4) bones without human activity traces. Taking into consideration the fact that cave bears and possibly also birds of prey did not cohabit the caves with humans, in order to determine the chronology of human occupation, we shall instead rely on radiocarbon dates provided by charcoal, bones with cut marks, or animal species which do not naturally live in caves, such as mammoths. If we take this into account, one can limit the list of reliable dating for LRJ into four measures i.e.:

- 33,100 ± 1200 BP (wood charcoal, Koziarnia Cave, layer F, Poz-110657, this study);
- 33,230 ± 480 BP (wood charcoal, Koziarnia Cave, layer H’, Poz-98901, this study);
- 32,500 ± 400 BP (woolly mammoth, Nietoperzowa Cave, layer 5b, Poz-23628 [32]);
- 38,160 ± 1250 BP (wood charcoal, Nietoperzowa Cave, layer 6, Gro-2181 [20]).

**Fig 14.**
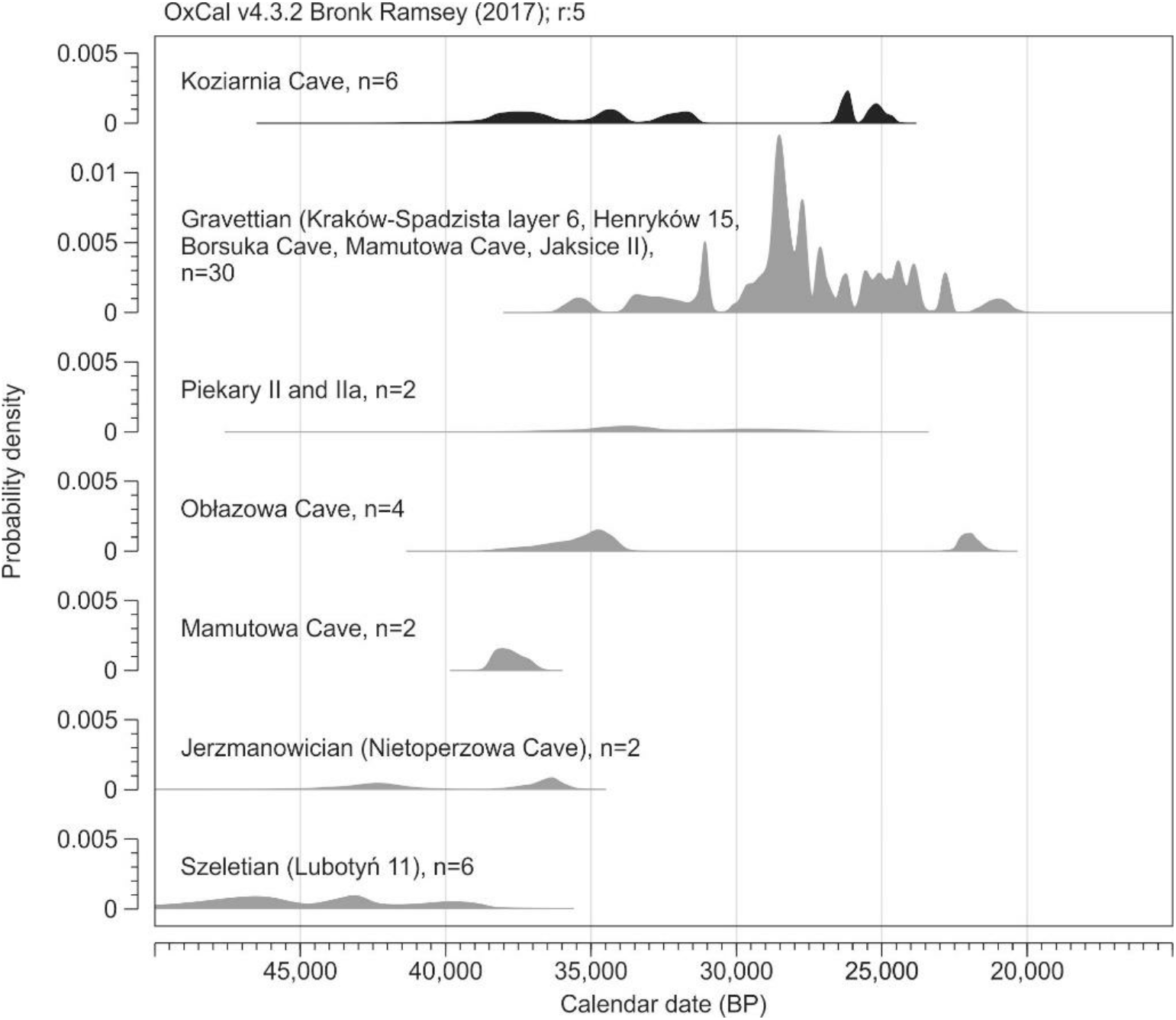
Correlation of the probability density distribution of radiocarbon dates from Koziarnia Cave, Middle-to-Upper Palaeolithic transition sites and early Upper Palaeolithic sites from southern Poland. Only the dates representing human settlement are regarded here (in the case of multi-strata cave sites – only the charcoal and reworked bones/ivory; in the case of single-stratum open-air sites – all the dates from a cultural layer). [Dates after 9, 20, 32, 86-98].

As long as the oldest date was obtained from the lower Jerzmanowician layer in Nietoperzowa Cave, one can assume that the presented set of dates demonstrates two separate settlement episodes. In such a case, the Jerzmanowician settlement in Koziarnia Cave could be tentatively correlated with the upper Jerzmanowician horizon in layer 4 in Nietoperzowa Cave, and represent the younger phase of Jerzmanowician.

Moreover, one should take into consideration the radiocarbon dates obtained recently on two bone points of the Mladeč type from Mamutowa Cave (38,5-36,5 ky cal. BP-S2), which overlap with those from Koziarnia Cave. Although Mladeč-type points are still believed to represent rather the Aurignacian tradition, in Mamutowa Cave no other Aurignacian artefacts were found either by Zawisza [84, 85] or Kowalski [76, 83]. Kowalski, who studied the stratigraphy in detail, determined a single Jerzmanowician layer (VI) and Gravettian occupation traces in loess layer 2. The radiocarbon-dated bone point comes from the old excavations by Zawisza; thus, their original stratigraphic position is impossible to establish, besides the information that they came from the inner part of the section.

The recent project on the ^14^C dating of the available Early Upper Palaeolithic bone points shows that they represent a broad chronology starting from 43 up to 34 ky cal. BP [94]. At several sites (Istalöskö, Dzerava Skala), such bone points were found in the company of leafpoints [99–104]. What is more, traces of Aurignacian occupation in Poland are very scarce and limited to several sites (e.g. Piekary II, Obłazowa layer VIII, Góra Puławska) [105–107], while the only available radiocarbon dates indicate the earliest Aurignacian settlement started around 36 ky cal. BP and is slightly younger than both bone points from Mamutowa Cave and the Jerzmanowician occupation in Koziarnia Cave.

The bone points from Mamutowa Cave are older than the oldest available Aurignacian dating north of the Carpathians, but at the same time they overlap with the Jerzmanowician settlement in Koziarnia and Nietoperzowa Caves. One can, therefore, assume that they could have originally belonged to the Jerzmanowician assemblage in Mamutowa Cave. Unfortunately, the chronology of Mamutowa Cave is mostly based on radiocarbon dates made on cave bear and bird bones. The sets of dates from underlying and overlying strata as well as from layer VI indicate some post-depositional sediment mixing (S2).

The possibility of the long chronology of the LRJ technocomplex exceeding the Campanian Ignimbrite (CI) eruption event is confirmed also by results obtained in Lincombian sites e.g. Beedings, indicating its lasting up to even 30 ky BP [108].

### Gravettian occupation

It is worth noting that aside from the Middle Palaeolithic and Jerzmanowician, also two Gravettian occupation phases can be identified in Koziarnia Cave. The first one could be correlated with the second and third human settlement phase recorded by the charcoal dated to 35-31 ky cal. BP (Fig 14). This dating overlaps with the earliest traces of the penetration of Gravettian hunters, confirmed recently in Henryków 15 in the Sudetes piedmonts. The chronology of the settlement traces in Koziarnia Cave indicates that the earliest Gravettian groups penetrated not only the nearest vicinities of the Moravian Gate but went much further into the Polish Highlands.

The second settlement phase, which is recorded only by two bone tools of uncertain stratigraphic positions, indicates Gravettian occupation in Koziarnia Cave simultaneous to a well-known mammoth butchering site at Spadzista Street in Kraków [74, 109,110]. Interestingly none of the Gravettian occupation phases in Koziarnia Cave overlap the Gravettian settlement from Mamutowa Cave, which can be dated to 29-27.5 ky cal. BP.

## Conclusions

The obtained results show the complex stratigraphy in Koziarnia Cave. Although the new fieldwork was conducted far from the cave entrance, where only scarce settlement traces could be found, detailed chronostratigraphic and archaeological analyses made possible to determine four general occupation phases in the cave. We may, therefore, assume that the cave was occupied in the Late Middle Palaeolithic, but the typo-technological character of the assemblage is still to be discussed. The site was also occupied during the Middle/Upper Palaeolithic transition. This occupation phase can be identified as Jerzmanowician and should be dated to 39-36 ky cal. BP. The obtained radiocarbon dates indicate that the Jerzmanowician tradition lasted longer and did not finish with the Campanian Ignimbrite eruption. Above the Jerzmanowician strata, a thin sterile layer can be observed, separating the overlying Gravettian strata. The earliest Gravettian occupation can be dated to 35-31 ky cal. BP, and thus represents the earliest Gravettian occupation in the Polish Jura, and one of the earliest to the North of Carpathians.

One should also emphasize that the recent results confirm the previous assumptions, claiming that humans and animals did not cohabit caves, even if their traces are found in the same lithostratigraphic layers. For this reason, only radiocarbon dates obtained either on charcoal fragments or modified bones or teeth should be used for determining human settlement at cave sites.

## Acknowledgements

Authors would like to thank Professor Teresa Madeyska, who give access to the archive field documentation of the site and provided a great support in our research. We would like to thank the Ojców National Park and the Institute of Geophysics of the Polish Academy of Sciences for their kind permission for conducting the fieldworks inside the cave. The ZooMS analysis was financed by the Max Planck Society and we would like to aknowledge Prof. Dr. Stefan Kalkhof and the IZI Fraunhofer institute of Leipzig for providing us access to a MALDI-TOF-MS.

## Supplements

**S1 File.** Stratigraphy of trench IX/2017 in Koziarnia.

**S2 Table.** Radiocarbon dates used in the probability density models.

**S3 Table.** Published radiocarbon dates of Jerzmanowician assemblages.

**S4 Table.** Morphometric data of stone artefacts.

**S5 File.** Traseological analysis of stone artefacts.

